# Butyrate blocks specific histone acetylation by preventing recruitment of p300 to acetylated histones

**DOI:** 10.1101/2025.10.10.681627

**Authors:** Muwei Jiang, Anthi Psoma, Jinxiao Lyu, Maria Gabriella Chiariello, Rutger Modderman, Frans Bianchi, Danny Incarnato, Siewert-Jan Marrink, Iris H. Jonkers, Geert van den Bogaart

## Abstract

Butyrate, a short-chain fatty acid (SCFA) produced by microbial fermentation of dietary fiber, exerts beneficial metabolic and immunomodulatory functions through hyperactivation of the histone acetylase (HAT) p300 and inhibition of histone deacetylases (HDAC). These effects are widely believed to result in the acetylation of distinct histones that regulate specific genes, including at transcription starting sites (TSS) and enhancers. However, we show that this is not the case, as butyrate dose-dependently increases the acetylation of histones throughout the entire genome. Molecular dynamics simulations suggest that the RING-loop prevents the recruitment of p300 to acetylated histones through its bromodomain, and thereby cannot maintain the acetylation of specific histones through positive feedback. This was confirmed by showing that only catalytically inactive p300 stably binds to acetylated histones. Thus, the epigenetic regulation of specific genes by butyrate is limited, but butyrate instead increases histone acetylation globally resulting in opening of the entire chromatin structure.

## Introduction

Short-chain fatty acids (SCFAs) are defined by the presence of an aliphatic tail of one to six carbon atoms, mainly produced from the intestinal microbial fermentation of dietary fiber in the gut. Acetate (C2), propionate (C3) and butyrate (C4) are the main SCFAs, which are produced in the human proximal colon in 70–140 mM concentration(Cummings et al., 1987). Butyrate exerts a variety of biological beneficial effects, including enhancing the intestinal epithelial barrier, maintaining intestinal mucosal immunity, anti-inflammation, and reducing IBD-associated symptoms(Canani et al., 2011; Liu et al., 2018). Although butyrate also signals through G-protein coupled receptors(Brown et al., 2003; Thangaraju et al., 2009) and PPAR-γ(Kinoshita et al., 2002; Alex et al., 2013), and can affect cellular metabolism and intracellular pH(Jiang et al., 2025a), the current scientific consensus is that butyrate exerts its physiological effects mainly by altering gene expression through epigenetic modifications related to increased histone acetylation. The acetylation of histones is a highly dynamic process regulated by two antagonistic families of enzymes: Histone acetyltransferases (HATs) use the substrate acetyl-CoA to covalently link the acetyl group to lysine residues, whereas histone deacetylases (HDACs) stabilize the chromatin structure by removing the acetyl group and restoring the side chain’s positive charge. Thereby, the concerted action of HATs and HDACs collectively orchestrates the dynamic regulation of gene expression through histone modification.

Historically, the increase in histone acetylation by butyrate has been largely attributed to the SCFA’s inhibitory effects on HDACs(Hamer et al., 2008) and many studies showed that butyrate increases histone acetylation in different cell types, presumably via HDAC inhibition (Davie, 2003; Sekhavat et al., 2007; Kim et al., 2009; Schulthess et al., 2019). For example, in human peripheral-blood monocyte-derived macrophages, butyrate enhances antimicrobial activity, allegedly through specific inhibition of HDAC3 as this phenotype can be copied by siRNA silencing and small molecule inhibitors of HDAC3(Schulthess et al., 2019).

More recently, butyrate was found to also result in hyperactivation of the HAT p300(Thomas and Denu, 2021). The catalytically activate HAT domain of p300 contains a lysine-rich autoinhibitory loop (AIL) that prevents binding to the histone substrate(Delvecchio et al., 2013). This AIL can become acetylated by another p300 molecule, resulting in the release of the AIL from the substrate binding pocket and the activation of p300(Delvecchio et al., 2013). Mass spectrometry showed that butyrate can also bind to the AIL, a process called butyrylation, also releasing it and resulting in hyperactivation(Thomas and Denu, 2021).

HDAC inhibition and HAT activation have been expected to result in acetylation of histones that regulate specific genes in macrophages(Schulthess et al., 2019), which are key immune cells of the intestinal tract(Zigmond and Jung, 2013). However, the precise histone acetylation sites that are affected by butyrate have not yet been mapped in macrophages. In this study, we therefore tested the effect of butyrate on histone acetylation in human monocyte-derived macrophages. Surprisingly, we found a reduction in specific histone acetylation sites by chromatin immunoprecipitation-sequencing (ChIP-seq), whereas microscopy and Western blot showed an increase in global histone acetylation. Molecular dynamics simulations suggested that AIL acetylation induces a conformational shift in p300, preventing its bromodomain from binding to acetylated histones. However, we could not confirm this conformational shift, as mutants predicted to disrupt it were either not expressed or only poorly expressed, potentially due to cytotoxicity. Moreover, the molecular dynamics simulations suggested that disruption of bromodomain binding to acetylated histones following AIL acetylation interferes with a positive feedback mechanism that is believed essential for maintaining specific histone acetylation sites(Thomas and Denu, 2021). We confirmed this mechanism by mutagenesis and small molecule inhibitors of p300, showing that only the catalytically inactive form of p300 is stably recruited to acetylated histones through its bromodomain. Thus, our data show that butyrate blocks specific histone acetylation by preventing recruitment of p300 to acetylated histones.

## Results

### Butyrate decreases specific histone acetylation at transcription start sites

Since butyrate results in a pronounced transcriptional reprogramming(Arpaia et al., 2013; Schulthess et al., 2019), we hypothesized that it might not only directly inhibit HDACs or activate HATs, but also affect their activity by altering their expression levels. We therefore determined the expression levels of the genes coding for HDACs and p300 in human monocyte-derived macrophages that were stimulated for 24 h with the pathogenic stimulus lipopolysaccharide (LPS), the inflammatory cytokine interferon (IFN)-γ, and a broad concentration range of butyrate (0.1–10 mM). This range is physiological as it is reflective of concentrations in different regions of the large intestine(Cummings et al., 1987) and these concentrations are also used in other *in vitro* studies(Boffa et al., 1978; Sealy and Chalkley, 1978; Davie, 2003; Chang et al., 2014; Jiang et al., 2025b, 2025a). mRNA levels of *HDAC3* were elevated upon increasing concentrations of butyrate, whereas *HDAC7* and *SIRT1* were downregulated and *HDAC1*, *HDAC4*, *HDAC8, SIRT2,* and *SIRT3* did not show a clear dose response (**Figure 1A** and **Supplementary Figure S1A**, **B**). Moreover, we also observed a dose-dependent reduction of *EP300* (the gene coding for p300) and induction of *HAT1* mRNA levels by butyrate (**Figure 1A** and **Supplementary Figure S1B**). Thus, butyrate differentially affects the expression levels of these HAT and HDACs, in line with widely accepted notion that butyrate regulates gene expression of specific genes.

**Figure 1.**
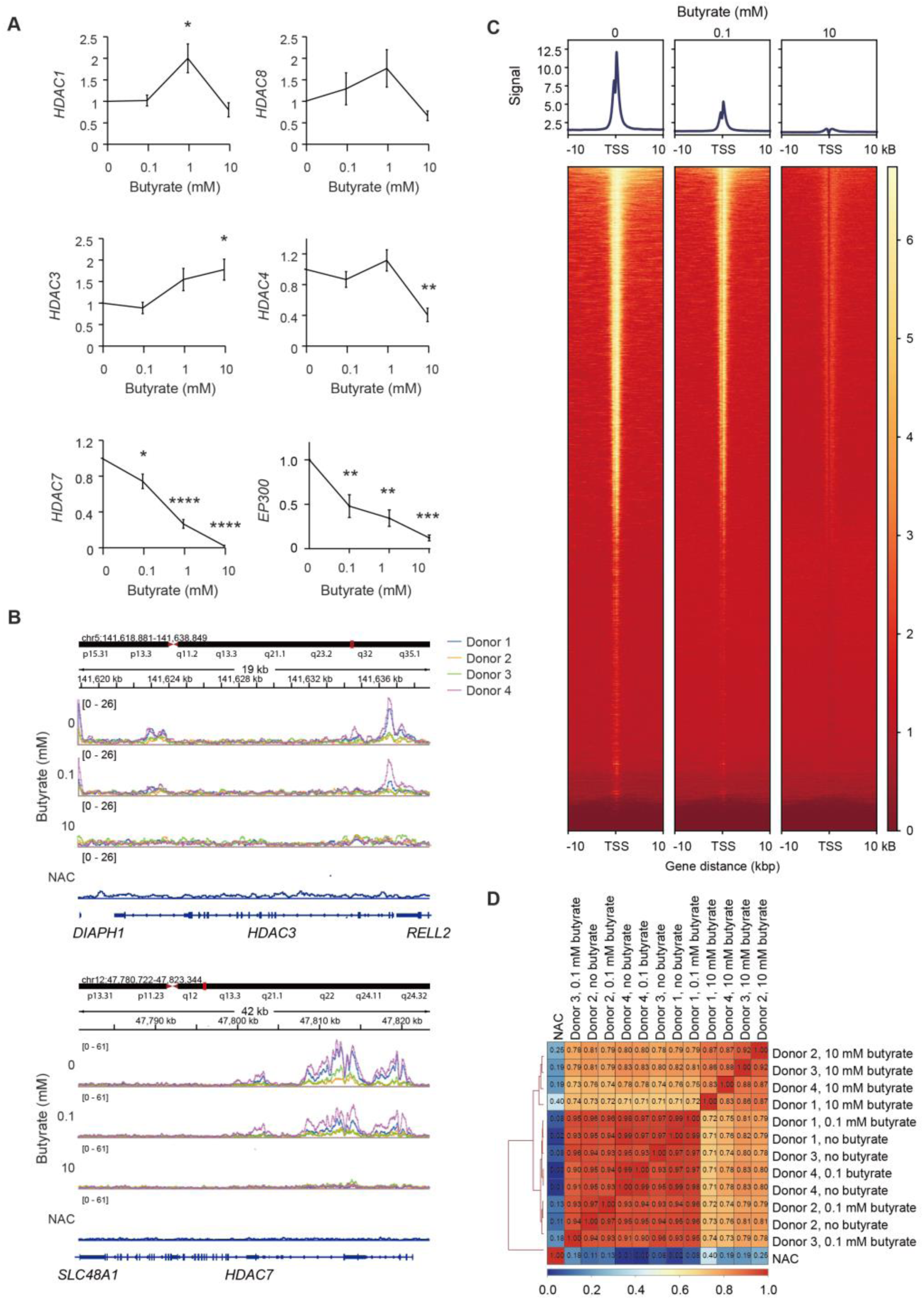
Butyrate decreases the enrichment of specific histone acetylation relative to total histone acetylation. Human peripheral blood mononuclear-derived macrophages were stimulated for 24 hr with LPS, IFN-γ and the indicated concentrations of Na-butyrate. (**A**) *HDAC* mRNA levels by RT-qPCR normalized to without butyrate. (One-way ANOVA with a Dunnett’s multiple comparisons test, n = 3 donors; error bars represent means ± SEM; *: P < 0.05; **: P < 0.01; ****: P < 0.0001; conditions were compared with the no butyrate condition) (**B**) Genome browser snapshots of published ChIP-seq data (Jiang et al., 2025b) showing H3K27Ac enrichment in *HDAC3, HDAC7* and flanking genes. NAC: no antibody control. (**C**) Intensity profile overlap of all transcription start sites (TSS), showing a dose-dependent decrease in fold-enrichment of peaks with Na-butyrate. (**D**) Correlation heatmap of the consensus peaks of the H3K27Ac signal in a window of -1 to 1 kbp around the 1000 TSS, showing that the 10 mM butyrate condition clusters together (n = 4 donors).

In addition to its effects on gene expression by epigenetic regulation, HDAC inhibitors such as TSA have been reported to post-transcriptionally regulate protein expression by decreasing mRNA stability(Xiong et al., 2005; Sharma et al., 2013). We therefore assessed whether the differential HDAC expression upon butyrate treatment correlated with histone acetylation at transcription starting sites (TSS) of HDAC encoding genes by a detailed analysis of our recent ChIP-seq data(Jiang et al., 2025b). We performed ChIP-seq on histone 3 K27-acetylation (H3K27Ac), as this is a canonical active enhancer mark associated with higher transcriptional activation(Delvecchio et al., 2013) and butyrate is well known to increase H3K27Ac(Schulthess et al., 2019). For these experiments, we used a polyclonal rabbit antibody raised against H3AcK27 (Diagenode, C15410196), because it has been used extensively in the literature for ChIP-seq. However, more recently it has been shown to also bind to other lysine acetylation sites of histones, which suggests that our findings could also apply to other histone acetylation sites.

Surprisingly, we observed that butyrate reduced the enrichment of H3K27Ac at the genes coding for all HDACs (relative to total reads), even for *HDAC3* despite it being higher expressed in the presence of butyrate (**Figure 1B** and **Supplementary Figure S1C**). This was also observed for the genes coding for calprotectin (*S100A8* and *S100A9*) that have been reported(Schulthess et al., 2019) to be strongly upregulated by butyrate in this cell type (**Supplementary Figure S1D**). In fact, this was a global phenomenon, as we consistently observed that butyrate dose-dependently reduced the enrichment of H3K27Ac at all transcription starting sites (TSS) (i.e. throughout the entire genome) for all four donors (**Figure 1C**, **D** and **Supplementary Figure S2A**). Butyrate also consistently reduced the intensity of common acetylation sites that were present in all four tested donors, identified by peak calling of H3K27ac for the conditions without butyrate (**Supplementary Figure S2B**). This reduction in histone acetylation by butyrate was also apparent when we identified regions with significant changes in acetylation levels using a sliding window of 1000 bp and DeSeq2 (**Supplementary Figure S2C**). Finally, histone acetylation was also consistently reduced by butyrate on enhancer regions (**Supplementary Figure S2D**). Because there is no data available on the locations of enhancer regions in monocyte-derived macrophages, we identified these as H3K4me1 positive, H3K27Ac positive, and H3K4me3 negative regions in their precursors, CD14-positive monocytes, using ENCODE data. Thus, contrary to our expectations, butyrate dose-dependently decreased H3K27 acetylation at specific sites relative to the entire genome.

### Butyrate increases histone acetylation globally

The ChIP-seq results showed that butyrate reduced the fold-enrichment of H3K27 acetylation peaks at specific sites relative to the entire genome. This seems contrary to the current model that butyrate increases histone acetylation(Hamer et al., 2008). However, this conclusion is largely based on bulk experiments which assess histone acetylation of the entire chromatin, mainly by incorporating radioactive acetate, altered electrophoretic mobility of histones, and Western blot analysis against acetylated histones(Boffa et al., 1978; Sealy and Chalkley, 1978; Davie, 2003). Therefore, the decrease of histone acetylation at specific sites that we observed by ChIP-seq might be caused by an increase in histone acetylation everywhere else, thus non-specifically throughout the entire chromatin. To confirm that butyrate increases the acetylation of total histones, we probed lysates of LPS and butyrate-treated macrophages by Western blot with the same antibody for H3K27 acetylation as used in our ChIP-seq (**Figure 2A**, **B** and **Supplementary Figure S3A**). We also probed the blot with an antibody recognizing multiple acetylation sites of H3 (K4, K9, K14, K18, K23, K27) (**Figure 2A**). Whereas 0.1 and 1 mM butyrate did not result in detectable levels of acetylated H3 by Western blot, 10 mM butyrate showed a clear increase in H3 acetylation, which is consistent with literature(Schulthess et al., 2019; Jiang et al., 2025a). Moreover, 10 mM butyrate also increased acetylation of H4 (K5, K8, K12, K16) (**Figure 4C**), showing that butyrate affects acetylation of multiple histones.

**Figure 2.**
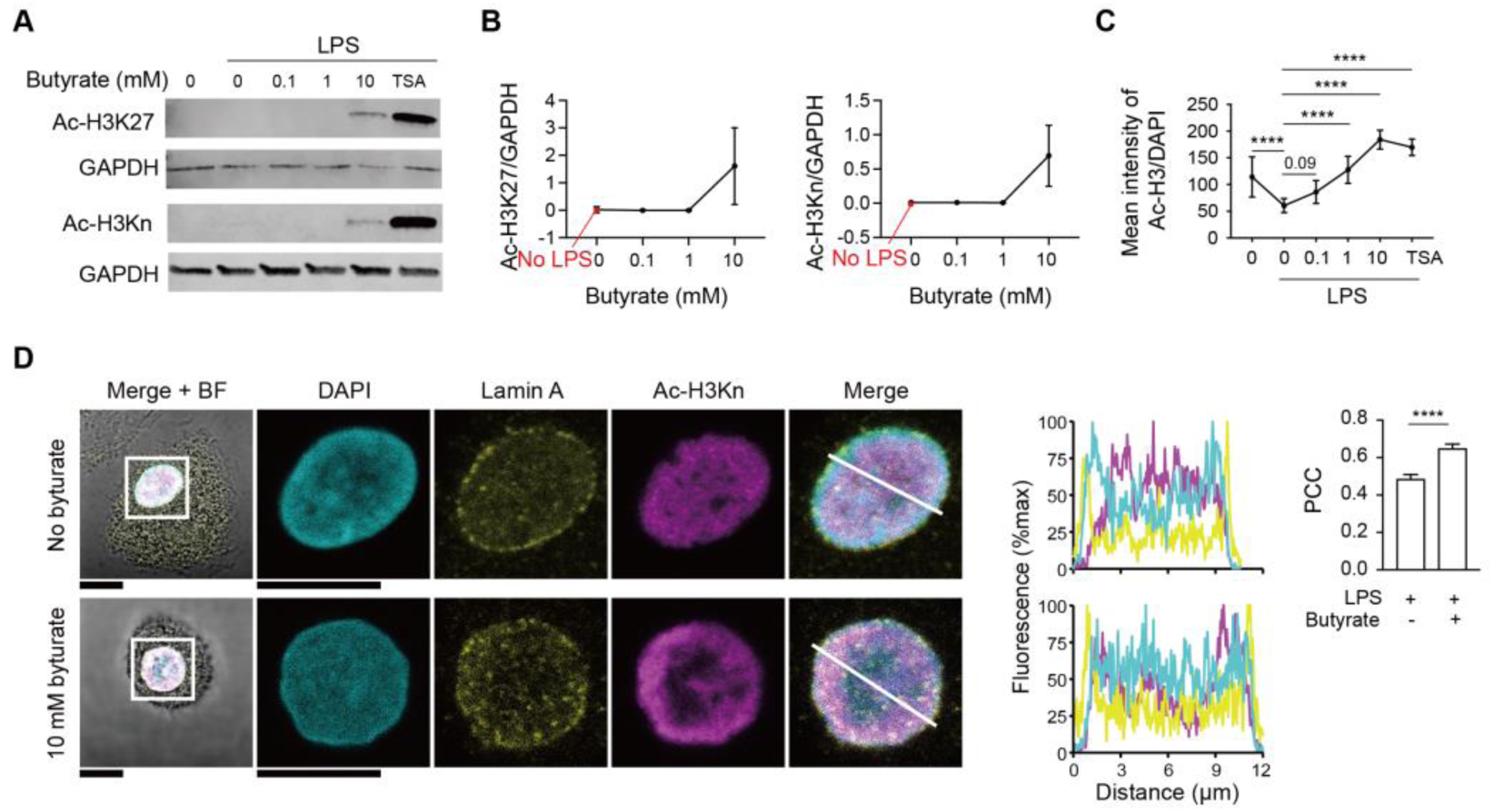
Butyrate induces hyperacetylation of histones globally. Human peripheral blood mononuclear-derived macrophages were stimulated for 24 hr with LPS, IFN-γ, and the indicated concentrations of Na-butyrate or 10 μM Trichostatin A (TSA). (**A**) Western blots showing histone acetylation detected with two different antibodies recognizing acetylated (Ac-)H3K27 and H3K4+9+14+18+23+27 (Ac-H3Kn) acetylation. (**B**) Quantification of Ac-H3K27 and Ac-H3Kn levels normalized to GAPDH. (n=3 donors; error bars represent means ± SEM; the complete blots for all donors are shown in Supplementary Figure S3A). (**C** and **D**), Immunofluorescence microscopy of macrophages labeled for Ac-H3Kn. (**C**) Quantification of the mean intensity of histone acetylation normalized to DAPI (One-way ANOVA with a Tukey’s multiple comparisons test, n = 3 donors, at least 5 cells per donor; error bars represent means ± SEM. ****: P < 0.0001; representative confocal images are shown in Supplementary Figure S3B). The macro for automated image analysis is provided in the Supplementary Data. (**D**) Representative confocal images showing immunolabeled nuclei for Ac-H3Kn (magenta), Lamin A (yellow), and DAPI (cyan). The line graphs show fluorescence intensity profiles indicated by the white line. The bar chart shows the Pearson correlation coefficient (PCC) between the Lamin A and H3Ac staining (paired 2-sided t-tests, n=3 donors, at least 5 cells per donor; error bars represent means ± SEM. Scale bars: 5 μm). The macro for automated image analysis is provided in the Supplementary Data.

To further address the discrepancy between the ChIP-seq and Western blot, we also quantified the levels of acetylated H3 by immunofluorescence microscopy. In line with our Western blot data, we observed a clear and dose-dependent increase of H3 acetylation in the nucleus (**Figure 2C**, **2D** and **Supplementary Figure S3B**). However, we noted that the distribution of acetylated histones differed markedly: Whereas acetylated H3 mainly located in foci that were distributed relatively uniformly throughout the nucleus in the absence of butyrate, it located more towards the edge of the nucleus in the presence of 10 mM butyrate (**Figure 2D**). This has been observed previously(Rada-Iglesias et al., 2007), and suggests that butyrate increases acetylation of heterochromatin, since heterochromatin is typically located at the edge of the nucleus and contains a higher density of histones than euchromatin. To quantify this phenotype, we co-stained for acetylated H3 and Lamin A, which is a major component of the nuclear lamina. Indeed, colocalization between these stainings was markedly increased in cells treated with 10 mM butyrate (**Figure 2D**). Therefore, we conclude that butyrate increases the acetylation of total histones throughout the entire chromatin, and thereby the specific acetylation of histones near TSS and at other specific sites is reduced relative to the rest of the chromatin.

### Activation prevents binding of p300’s bromodomain to acetylated histones

Next, we explored the mechanism of how butyrate results in loss of acetylation of specific histones, but increases histone acetylation overall. We focused on p300, because, despite the decreased expression of p300 (**Figure 1A**), butyrate has been shown to hyperactivate p300(Thomas and Denu, 2021) and the global increase in hyperacetylation is suggestive of a role for hyperactivated p300 instead of selective inhibition of specific HDACs. Moreover, H3K27 and other lysines on H3, H4, and other histones are main sites for p300 acetylation(Henry et al., 2013), and butyrate-induced hyperacetylation of H3K27 has been shown to be blocked by the p300 inhibitor A485(Thomas and Denu, 2021). The catalytically active core of p300 consists of a bromodomain, a plant homeodomain (PHD) with the inhibitory RING loop, and the HAT domain(Delvecchio et al., 2013). p300 acetylates histones at specific enhancer regions by being recruited to active transcription factors(Ortega et al., 2018). In addition, p300 binds to acetylated histones with its bromodomain and this also contributes to H3K27 acetylation(Raisner et al., 2018), likely because this substrate targeting of p300 aids in the maintenance of acetylated histone peaks by positive feedback(Delvecchio et al., 2013).

Comparison of a cryo-EM structure of the p300 catalytic core bound to the acetylated H4 nucleosome(Kikuchi et al., 2023) with a cryo-EM structure of free p300(Ghosh et al., 2019) suggests that the inhibitory RING-loop undergoes a conformational change and moves between the HAT and the bromodomains upon activation (**Supplementary Figure S4**). Moreover, not only lysine residues located in the AIL are autoacetylated, but also lysines in the RING-loop of the PHD domain that connects the bromodomain with the HAT(Black et al., 2008). This acetylation might affect the interactions of the RING-loop with the HAT and bromodomain. We therefore hypothesized that acetylation or butyrylation of the AIL would promote the interaction of the RING-loop with the bromodomain, thereby preventing its binding to acetylated histone substrates.

We conducted molecular dynamics (MD) simulations of p300 to test this hypothesis. The MD simulations were based on a reference structure (PDB: 6GYR)(Ortega et al., 2018) of p300 containing the bromodomain, PHD-domain (including RING-loop) and HAT (residues 1045–1664). As we were specifically interested in the interactions between the bromodomain and RING-loop upon acetylation, we performed the first set of simulations on a smaller system composed of the bromodomain and PHD-domain (**Figure 3A**, **B**, and **Supplementary Figure S5**). Besides the non-acetylated form, we simulated the triple acetylated form K1180Ac, K1203Ac, and K1228Ac, with three lysines in the RING-loop that have been shown to be acetylated(Black et al., 2008) (see Methods section for computational details).

**Figure 3.**
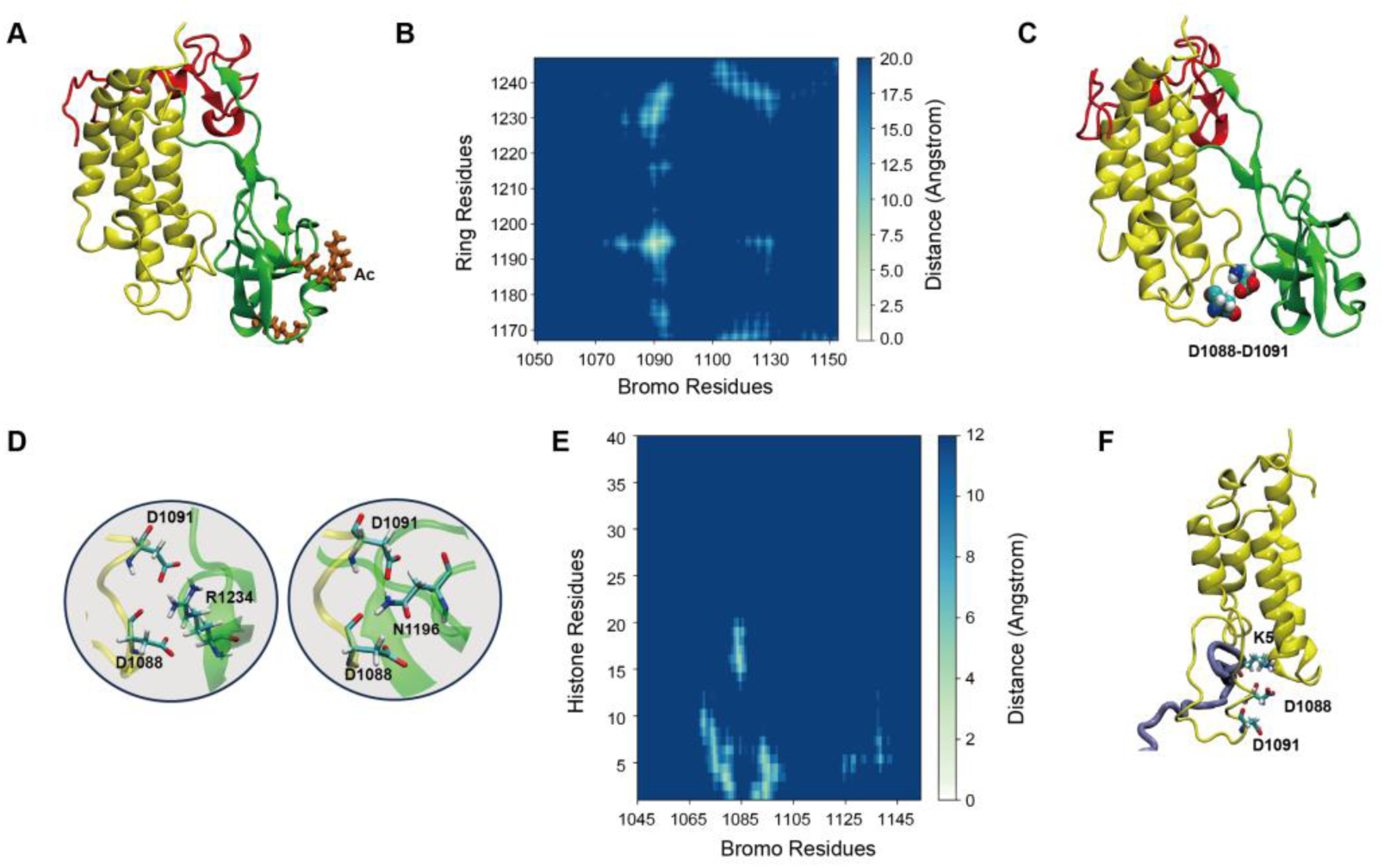
MD simulations of the p300 acetyltransferase system. The reference structure is PDB 6GYR (**A**-**D**). Different protein sub-units are color-coded: bromodomain (yellow), RING-loop (green), and PHD domain (red). (**A**) Structure extracted from the simulation of the triple mutant K1180Ac, K1203Ac, and K1228Ac, with acetylated lysines highlighted in orange. The bromodomain and RING-loop are in contact. (**B**) Contact map between the RING-loop and bromodomain residues, averaged over the last 500 ns of simulation. (**C**) Structure detailing the interaction between the bromodomain and RING-loop, showing that residues D1088 and D1091 are primarily involved in binding. (**D**) Interaction between residues D1088 and D1091 (bromodomain) with N1196 and R1234 (RING-loop). (**E**-**F**) MD simulations of p300 with the bromodomain bound to the N-terminal chain of histone H4 (ice blue). The reference structure is PDB 8HAG. The contact map between the bromodomain and histone protein chain residues is shown, along with a representative structure extracted from the simulation.

Based on a 1 μs simulation we generated a contact map between the bromodomain and RING-loop. The bromodomain interacted stably with the RING-loop when the lysines were acetylated, while this interaction did not take place in the non-acetylated system (**Supplementary Figure S5**). However, this interaction is not directly mediated by the acetylated lysines. Instead, simulations of the complete non-acetylated system (**Supplementary Figure S6**) revealed that the charged lysines and aspartates participate in a network of interactions between the RING-loop and HAT. When these lysines are neutralized by acetylation, these interactions are disrupted, allowing the RING-loop to detach from the HAT and move closer to the bromodomain.

Focusing on the interactions between the bromodomain and RING-loop, we found that this involves a specific segment of the bromodomain (loop 1080–1095). Notably, two aspartate residues (D1088 and D1091) in the bromodomain contact arginine and asparagine residues in the RING-loop (N1196 and R1234), as shown in representative structures and contact plots over time from the simulation (**Figure 3C**, **D** and **Supplementary Figure S7**). These aspartates are located on a highly flexible loop (see root mean square fluctuation (RMSF) in **Supplementary Figure S7A**), which adapts to form the binding region for the histone peptide chain. The interaction of D1088 and D1091 with the RING-loop thus could compete with binding of acetylated histones.

In the structure of p300 bound to the nucleosome (PDB: 8HAG)(Kikuchi et al., 2023) the eleven N-terminal residues of H4 remain unresolved. To clarify whether residues D1088 and D1091 of the bromodomain interact with the histone peptide, we reconstructed the system using the reference structure 8HAG. Here, the peptide chain of H4 containing the eleven missing N-terminal residues was rebuilt. **Figure 3E** and **F** show a contact map between the bromodomain and the first 40 residues of the histone peptide chain, along with a representative structure illustrating this interaction. The first 15 residues of the peptide are stably bound to the bromodomain, with acetylated lysine 12 buried in the bromodomain. The preceding residues wrap around the bromodomain, including the region spanning residues 1090–1100. This revealed the interaction between K5 and D1091 (**Supplementary Figure S8**).

In conclusion, the MD simulations indicate that the bromodomain interacts with either the RING-loop or the histone peptide chain using the same residues, suggesting that binding to one excludes binding to the other. These results thus suggest that if the bromodomain and RING-loop contact through aspartates D1088 and D1091 of the bromodomain, this prevents binding with the histone chain. Alternatively, as the RING-loop approaches the bromodomain, the resulting interactions could weaken the binding with the histone peptide, causing it to detach. The binding of the bromodomain to acetylated histones is believed to allow for the maintenance of specific histone acetylation sites by positive feedback(Delvecchio et al., 2013). As butyrate results in butyrylation of p300’s AIL(Thomas and Denu, 2021), the loss of specific histone acetylation peaks in butyrate-treated cells might be caused by the loss of p300 binding to acetylated histones.

### Butyrylated p300 hyperacetylates histones globally

We tested the prediction from the MD simulations that p300 with acetylated AIL would show reduced binding of its bromodomain to acetylated histones. We tested this by assessing the effect of butyrate on the binding of p300 to acetylated H3 using immunoprecipitation experiments. As the levels of endogenous p300 in human monocyte-derived macrophages were too low for immunoprecipitation experiments and these cells are difficult to transfect with high efficiency, we overexpressed GFP-tagged residues 1048–1664 of p300, consisting of its bromodomain, PHD with RING-loop, and HAT domain, in HeLa cells(Schneider et al., 2022). Western blot on complete cell lysates (inputs) showed that 10 mM butyrate increased H3 and H4 acetylation in the HeLa cells, which could be partially inhibited by the HAT inhibitor A485 that binds to the catalytic site of the HAT domain(Lasko et al., 2017), but not by the bromodomain inhibitor ICBP-112 (**Figure 4A** and **Supplementary Figure S9A-D**). A485 treatment did not completely block butyrate-induced histone acetylation, likely because of the activity of other HATs and butyrate’s inhibition of HDACs.

**Figure 4.**
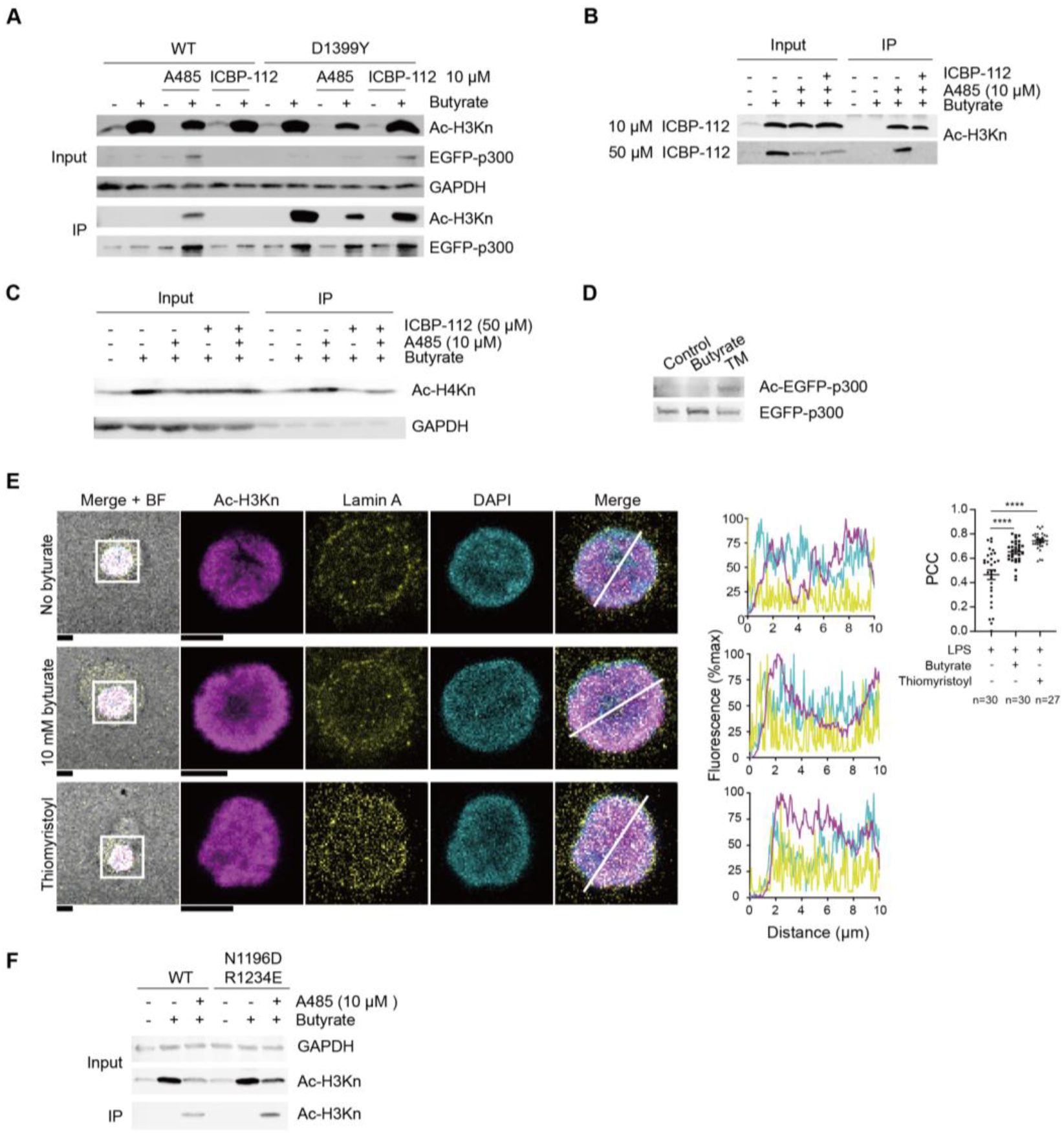
Butyrylated p300 does not bind to acetylated H3. (**A**) Hela cells were transfected with the EGFP tagged catalytic core of p300 wildtype or catalytically inactive D1399Y mutant, and stimulated for 24 hr with 10 mM Na-butyrate, 10 μM HAT inhibitor A485, and/or 10 μM bromodomain inhibitor ICBP-112 prior to immunoprecipitation (IP) for EGFP. Western blot showing Ac-H3K4+9+14+18+23+27 (Ac-H3Kn), EGFP-p300, and GAPDH detected in lysates and IP samples. (**B**) Transfected Hela cells were stimulated for 24 hr with 10 mM Na-butyrate, 10 μM A485, and 10 or 50 μM ICBP-112, showing that A485-induced binding of p300 to acetylated histones could be blocked by 50 μM ICBP-112. (**C**) Transfected Hela cells were stimulated for 24 hr with 10 mM Na-butyrate, 10 μM A485, and 50 μM ICBP-112. Western blot showing Ac-H4K5+8+12+16 (Ac-H4Kn) in lysates and IP samples. (**D**) Transfected Hela cells were stimulated for 24 hr with 10 mM Na-butyrate and 10 μM of SIRT2 inhibitor Thiomyristoyl. Western blot probed with an antibody recognizing an acetylated lysine located in the AIL of p300 (K1542), showing that acetylated p300 was increased by TM treatment. (**E**) Human peripheral blood mononuclear-derived macrophages were stimulated for 24 hr with LPS, IFN-γ and 10 mM Na-butyrate or 10 μM TM. Representative confocal images showing immunolabeled nuclei for Ac-H3Kn (magenta), Lamin A (yellow), and DAPI (cyan). Scale bars: 5 μm. The line graphs show fluorescence intensity profiles indicated by the white line. The Pearson correlation coefficient (PCC) between the Lamin A and Ac-H3Kn staining (One-way ANOVA with a Dunnett’s multiple comparisons test, 3 donors were analyzed; >8 cells per donor (exact counts in figure); error bars represent means ± SEM; ****: P < 0.0001. More representative cells are shown in Supplementary Figure S9H). The macro for automated image analysis is provided in the Supplementary Data. (**F**) Hela cells were transfected with N1196D and R1234E mutant, and stimulated for 24 hr with 10 mM Na-butyrate, 10 μM HAT inhibitor A485 followed by IP for EGFP. All complete blots and quantifications of 3 independent experiments are shown in Supplementary Figure S9 and 10.

In line with the predictions from the MD simulations, A485 treatment resulted in stable binding of p300 to acetylated H3 and H4 histones as shown by the immunoprecipitation (**Figure 4A-C** and **Supplementary Figure S9A-H**). A catalytically inactive mutant of p300, carrying the canonical D1399Y mutation located within the catalytical center of the HAT domain(Delvecchio et al., 2013), also bound stably to acetylated H3. This binding was reduced by 50 μM ICBP-112 (**Figure 4B** and **Supplementary Figure S9D-F**). Similar results were observed for acetylated H4 (**Figure 4C** and **Supplementary Figure S9G-H**). Thus, only catalytically inactive p300 stably binds to acetylated histones via its bromodomain, whereas activated p300 (i.e. with acetylated or butyrylated AIL) does not.

To confirm that the hyperactivation of p300 results in loss of specific histone acetylation, we performed experiments with Thiomyristoyl, a small molecule inhibitor of Sirtuin 2 (SIRT2). SIRT2 is a class III HDAC that deactivates p300 by deacetylating its AIL(Black et al., 2008). While other HDACs can also deacetylate p300, SIRT2 plays a key role in this process, as its inhibition by Thiomyristoyl has been shown to increase AIL acetylation(Black et al., 2008). This was confirmed by Western blot using an antibody specific for acetylated K1542 in the AIL of p300, which demonstrated that Thiomyristoyl treatment enhanced AIL acetylation in overexpressed p300 in HeLa cells (**Figure 4D** and **Supplementary Figure S9I**). Moreover, immunofluorescence microscopy of human monocyte-derived macrophages showed that acetylated H3 located more strongly toward the nuclear rim upon Thiomyristoyl treatment, thus phenocopying the effect of butyrate treatment (**Figure 4E** and **Supplementary Figure S9J**). These findings strengthen our conclusion that p300 with acetylated or butyrylated AIL is not recruited to acetylated histones.

To test the prediction from the MD simulation that the interaction of D1088 and D1091 with the RING-loop competes with binding of acetylated histones, we mutated the N1196 and R1234 residues of the RING-loop and performed immunoprecipitation experiments on Hela cells. Conversion of only the N1196 into an aspartic acid had no effect. However, as predicted by our molecular dynamics simulations, Western blot results showed that in the condition with co-treatment with butyrate and A485, the p300 double mutant N1196D and R1234E bound stronger to acetylated histones compared to wildtype (**Figure 4F** and **Supplementary Figure S10**). We also tested the D1088R and D1091R double mutant of p300, as we expected these mutations to enhance binding to acetylated histones even further. However, this mutant did not express, potentially because of cellular toxicity or misfolding. Similarly, p300 with K1180, K1203, and K1228 converted to arginine, which cannot be acetylated and therefore was predicted to result in stable binding of the bromodomain to acetylated histones, was only expressed at very low levels suggestive of cytotoxicity.

Together, our findings show that butyrate results in loss of acetylation of specific histones, but increases histone acetylation overall, in line with previous findings(Rada-Iglesias et al., 2007). Mechanistically, our MD simulations and immunoprecipitation experiments show that only inactive p300 stably binds to acetylated histones through its bromodomain. The activation of p300 by acetylation or butyrylation results in dissociation of the bromodomain from acetylated histones. Since the recruitment of p300 to acetylated histones is believed to allow for the maintenance of specific histone acetylation sites by positive feedback(Delvecchio et al., 2013), the loss of specific histone acetylation peaks in butyrate-treated cells are likely caused by the loss of p300 binding to acetylated histones.

In the final experiment, we tested the functional consequences of p300 inhibition of the production of the canonical inflammatory cytokine TNF-α by macrophages that were stimulated with LPS and IFN-γ. p300 blockage by A485 or ICBP-112 in the absence of butyrate resulted in an increase of TNF-α production, although this effect was not significant with the inhibitors alone **(Supplementary Figure S11)**. However, in the presence of 10 mM butyrate, we observed the opposite effect and p300 blockage by A485, ICBP-112 or both compounds together almost completely blocked TNF-α production **(Supplementary Figure S11)**.

## Discussion

Butyrate is well known to exert beneficial biological functions by promoting histone acetylation at promoter and enhancer regions of specific genes(Hamer et al., 2008; Canani et al., 2011; Liu et al., 2018). However, our data now show that this is not the case: butyrate-mediated histone acetylation is not site-specific, but instead dose-dependently increases histone acetylation globally over the entire chromatin.

A comparison of published cryo-EM structures of the p300 catalytic core bound to acetylated H4(Kikuchi et al., 2023) and free p300(Ghosh et al., 2019) suggested that the inhibitory RING-loop of the PHD-domain can move between the HAT and bromodomain upon activation (**Supplementary Figure S4**). Indeed, our MD simulations support a model where acetylation of lysine residues located on the RING motif weaken the interaction of the RING-loop with the HAT and consequently the RING-loop moves closed to the bromodomain (**Supplementary Figure S5** and **S6**). Here, a conserved asparagine and arginine located on the RING-loop interact with conserved aspartic acids of the bromodomain, causing the histone to no longer bind to the bromodomain (**Figure 3B**, **E** and **F**). Therefore, we predicted that upon butyrylation of the AIL, the bromodomain of p300 can no longer bind to acetylated histones as its binding site is occupied by the RING-loop. However, since the p300 mutants D1088R, D1091R, K1180R, K1203R, and K1228R, predicted to disrupt RING-loop interactions with the bromodomain and thereby stabilize its interaction with acetylated histones, were either not expressed or expressed only at very low levels, we were unable to experimentally confirm the proposed conformational change, possibly due to mutant-induced cytotoxicity. Thus, further studies are required to experimentally validate the consequences of RING-loop acylation on p300 conformation.

Nevertheless, we confirmed the predictions from the molecular dynamics simulations that only inactive p300 without acetylated AIL is recruited to acetylated histones via its bromodomain using the p300 inhibitor A485 and the catalytically inactive D1399Y mutant, which cannot be auto-acetylated (Delvecchio et al., 2013) (**Figure 4A-C**). As the binding of multiple p300 molecules to chromatin is well known to result in p300 activation by trans-acetylation of its AIL (Ortega et al., 2018), our findings suggest that inactive p300 is recruited to acetylated histones and subsequently dissociates upon its activation (**Figure 5**). This dissociation of active p300 likely allows for the acetylation of other histones nearby, and thereby for maintenance of acetylated histone sites through positive feedback (Delvecchio et al., 2013). However, this feedback cannot occur in the presence of butyrate, since butyrate activates p300 by butyrylation of its AIL (Thomas and Denu, 2021) and our data show that butyrylated p300 is not recruited to acetylated histones (**Figure 4A**). We therefore propose that butyrylated p300 acetylates histones at random, resulting in the global hyperacetylation observed by Western blot and microscopy (**Supplementary Figure S12**). Although we showed that butyrate significantly decreased *EP300* mRNA expression in human macrophages, our Western blot results showed the clear presence of endogenous K1542 acetylated p300 (**Supplementary Figure S9I**).

**Figure 5.**
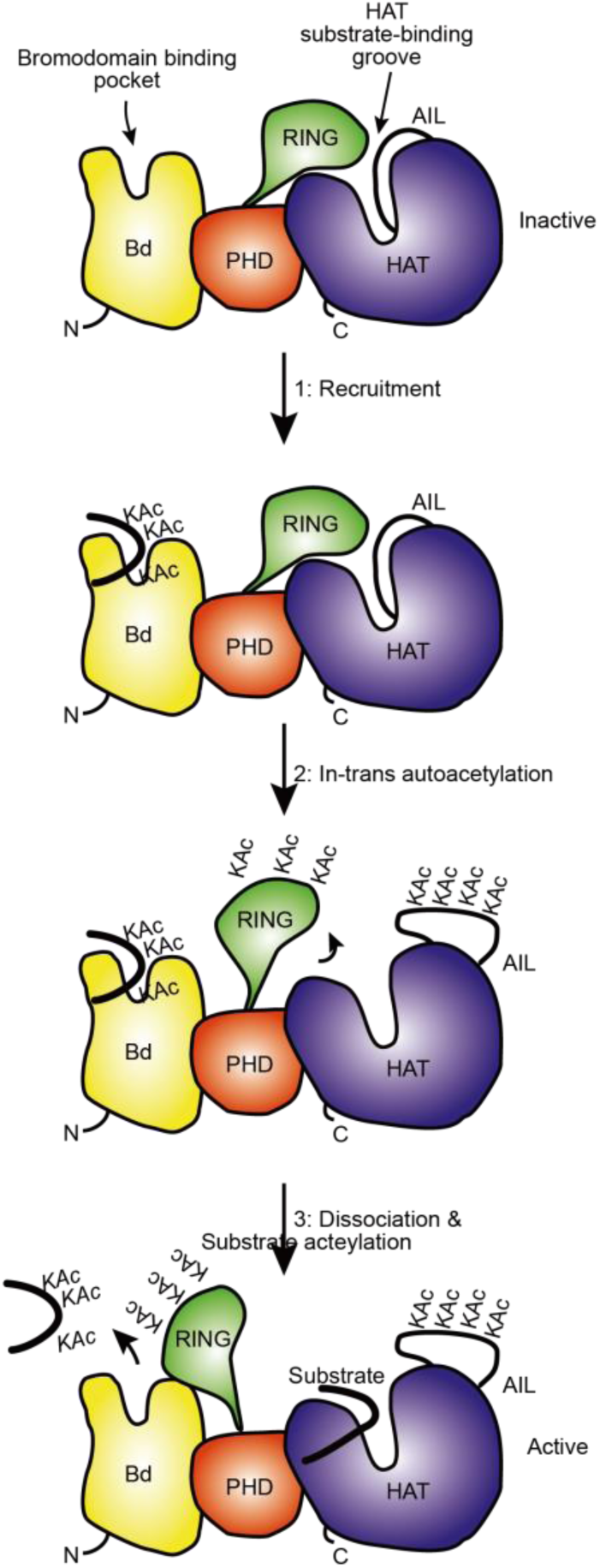
Model for p300 regulation and substrate acetylation. In the inactive state (top), the RING loops blocks the catalytic site of the HAT domain (Liu et al., 2008). The bromodomain of p300 is recruited to acetylated histones (step 1). As proposed previously, the recruitment of at least two copies of p300 result in the in-*trans* autoacetylation of the autoinhibitory loop (AIL) and displacement of the AIL and RING-loop, thereby exposing the catalytic site of the HAT domain and activating substrate acetylation (step 2) (Delvecchio et al., 2013). Lysine residues on the RING-loop also become autoacetylated (Black et al., 2008), which weakens interactions between the RING-loop and the HAT. The RING-loop now interacts with the bromodomain, causing its dissociation from acetylated histones (step 3). Yellow: bromodomain (Bd); green: RING-loop; red: PHD-domain; blue: HAT; KAc: acetyl-lysine. Model adapted from (Delvecchio et al., 2013).

Indeed, we showed that the inhibition of SIRT2 with Thiomyristoyl, which increases the acetylation of the AIL of p300 (**Figure 4D**), phenocopies the effect of butyrate treatment and results in a similar distribution of histone acetylation (i.e., at heterochromatin). Moreover, since *in vitro* activity assays showed that SIRT2 is about 4-fold less active towards butyrylated than acetylated substrates (Feldman et al., 2013), SIRT2 will likely less efficiently deactivate butyrylated p300 which could further increase the global histone hyperacetylation.

Our study contrasts many, but not all (Rada-Iglesias et al., 2007), studies where it is found by ChIP-qPCR and ChIP-seq (Furusawa et al., 2013) that butyrate increases histone acetylation at specific promoter sites. These differences are unlikely to be the result of different butyrate concentrations, as we already observed the increase in global histone acetylation at 0.1 mM butyrate by microscopy, which is lower than used in most studies, and the effect is highly pronounced at 10 mM, which is higher than used in most studies. Notably, this increase in histone acetylation was not detected by Western blot, likely due to insufficient sensitivity. Although we cannot rule out cell type specific effects, we believe our mechanism is general given the key role of p300 (and its close paralog CBP) in most cell types. Moreover, ChIP-qPCR might be prone to two experimental artefacts. First, ChIP-qPCR data is usually normalized against the input, but as we show that butyrate increases acetylation of histones globally, this might result in higher levels of immunoprecipitated genes, especially with low DNA fragmentation. Second, ChIP data is sometimes normalized against other genes that are not expected to be altered upon butyrate treatment, like *RPL13*, *CKS2* and *GAPDH*. However, butyrate also affects histone acetylation at those genes (**Supplementary Figure S1E**). In line with this, our ChIP-qPCR showed a reduction in histone acetylation of *CKS2* and *GAPDH* by 0.1 and 10 mM butyrate (**Supplementary Figure S1E** and **F**).

A question that is raised by our study is how butyrate can result in both up-and downregulation of specific genes, especially as it has been shown by pertussis toxin and overexpression that this is independent to G-protein coupled receptor (GPCR) signaling in macrophages (Chang et al., 2014; Schulthess et al., 2019). In addition to its effects on HATs, HDACs and GPCRs, butyrate has been reported to activate the peroxisome proliferator-activated receptor gamma (PPAR-γ), a nuclear receptor that functions as a transcription factor and can regulate the expression of multiple genes (Kinoshita et al., 2002; Alex et al., 2013). Moreover, especially at higher concentrations, butyrate promotes fatty acid oxidation and thus alters metabolic signaling cascades (Liu et al., 2018). We recently showed that at acidic extracellular pH, butyrate is also accumulated within cells and can acidify the cells(Jiang et al., 2025a). The beneficial biological functions exerted by butyrate can likely be attributed to the combination of global histone acetylation, GPCR and PPAR signaling, and metabolic and pH effects.

Consistent with the idea that butyrate acts through multiple mechanisms, we recently demonstrated that its effects on macrophages are highly concentration-dependent. Specifically, 0.1 mM butyrate suppresses LPS-induced TNF-α production via a mechanism involving PPAR-γ signaling, whereas 10 mM butyrate promotes IL-1β production and reduces anti-inflammatory IL-10 levels through pathways involving G protein-coupled receptors, the lipid transporter CD36, and the kinase SRC (Jiang et al., 2025). Although still a hypothesis, this dual effect of butyrate may explain the contrasting impact of p300 inhibition on TNF-α production by macrophages in the presence versus absence of butyrate (**Supplementary Figure 11**). It is possible that the pro-inflammatory effects of high butyrate concentrations require p300 hyperactivation, while at low concentrations, p300 plays a more anti-inflammatory role. In any case, our data underscore the complexity of butyrate-mediated regulation of macrophage-driven inflammation, which cannot be attributed solely to epigenetic control of specific genes via histone acetylation.

## Materials and methods

### Cell lines and cell culture

HeLa cells were cultured in Dulbecco’s Modified Eagle’s Medium (DMEM, Gibco) supplemented with 10% fetal bovine serum (FBS, Thermo Fisher Scientific), 1% antibiotic-antimitotic solution (AA, Gibco), and 1% L-glutamine (Gibco) at 37 ℃ and 5% CO_2_. Peripheral blood mononuclear cells (PBMC) were obtained from buffy coats of human blood donors by ficoll gradient centrifugation. Buffy coats were obtained anonymized from the Dutch blood bank. All blood donors were informed about the research and granted their consent to the blood bank. The research is exempt from ethical approval by the Dutch Medical Research with Human Subjects Law, because samples were provided fully anonymized to the researchers and the blood donors did not have to undergo extra procedures. CD14^+^ monocytes were isolated from PBMC using CD14 MicroBeads (Miltenyi). Human peripheral blood mononuclear-derived macrophages were generated in Ultra-low adherent 6-well plates (Corning) in 2 mL RPMI 1640 medium (Gibco) supplemented 10% FBS, 1% antibiotic-antimitotic solution (AA, Gibco), and 1% L-glutamine (Gibco) and M-CSF (RnD 216-MC, 100 ng/mL) at 37°C and 5% CO_2_ for seven days. On day 4, macrophages were supplemented with 1 mL medium containing 50 ng/mL M-CSF.

### Reverse transcription–quantitative PCR (RT-qPCR)

1×10^6^ human peripheral blood mononuclear-derived macrophages per condition were treated with 1 μg/mL LPS, 20 ng/mL IFN-γ and 0.1 mM, 1 mM or 10 mM sodium butyrate (Sigma) for 24 hours. Cells were lysed and then RNA was isolated from the lysate using Quick-RNA Miniprep Kit (ZYMO), cDNA was synthesized with random hexamer primers (Roche) and the M-MLV Reverse Transcriptase kit (ThermoFisher). RT-qPCR was performed using 4 ng of cDNA with PowerUP SYBR Green Master Mix, and the 10 μL for each reaction was incubated at 50°C for 2 min, 95°C for 2 min, followed by 40 cycles of denaturation (95°C for 15 sec), annealing and extension (60°C for 1 min) with a BioRad CFX96 qPCR System. The primer sequences are listed in Supplementary Table 1. Analyzed genes were normalized to Small Nuclear Ribonucleoprotein D3 Polypeptide (SNRPD3).

### ChIP-seq

The ChIP-seq wperformed as described (Jiang et al., 2025b)10/10/2025 12:51:00 PM. 1×10^6^ human peripheral blood mononuclear-derived macrophages per condition were treated with 1 μg/mL LPS, 20 ng/mL IFN-γ and 0.1 mM or 10 mM sodium butyrate for 24 hours. Chromatin preparation and library construction were carried out using on the standard 2014 Blueprint Histone ChIP and Kapa Hyper Prep kit protocol (http://www.blueprint-epigenome.eu) and the reagents described therein. For chromatin preparation, we established the optimal fixation time to be 5 minutes in 1% formaldehyde at room temperature. The shearing time was 10 or 12 minutes using the Covaris Focused-ultrasonicator (S220) and the low cell protocol for chromatin shearing, which is optimized for chromatin shearing of 1 to 3 million cells using 13 icrotubeTUBE-130 with AFA fiber screw cap (Covaris, cat 520216). Immunoprecipitation was carried out with polyclonal rabbit H3AcK27 antibody (Diagenode, C15410196). After the library construction, samples were PAGE-purified, quantified on the Agilent tapestation 2200 bioanalyzer, and equimolarly pooled. Sequencing was done on the DNBseq platform using Paired-End 100 sequencing.

### Analysis of ChIP-seq data

ChIP-seq data was aligned to the hg38 assembly of the human genome using Bowtie v1.3.1 (parameters: -l 28 -n 2 -–1 --b–t --str–a --chunkmbs 3200) (Langmead et al., 2009), retaining only uniquely aligned reads. Correlation, principal component analysis, heatmap generation were performed using deepTools v3.5.0 (Ramírez et al., 2016).

### RNA-seq

PBMC-differentiated macrophages were seeded in 6-well plate at 1,000,000 cells/well, cells were treated with 1 μg/mL LPS, 20 ng/mL IFN-γ and 0.1 mM, 1 mM or 10 mM sodium butyrate for 24 hours. Samples were washed twice in PBS and lysed by direct addition of 1 mL TRIzol reagent (ThermoFisher Scientific, 15596026). RNA extraction was performed as per manufacturer instructions, and RNA integrity was assessed by electrophoresis. Library preparation was performed using the NEBNext Ultra II RNA Library Prep Kit for Illumina (New England Biolabs, E7770L), as per manufacturer instructions. Libraries were sequenced on an Illumina NextSeq 1000 sequencer.

RNA-seq data was aligned to the hg38 assembly of the human genome using STAR v2.7.10b (parameters: --outFilterMultimapNmax 20 --alignSJoverhangMin 8 --alignSJDBoverhangMin 1 --outFilterMismatchNmax 999 -- outFilterMismatchNoverReadLmax 0.04 --alignIntronMin 20 --alignIntronMax 1000000 --alignMatesGapMax 1000000). Gene counts were computed using featureCounts v2.0.0 and the latest refFlat annotation from UCSC (parameters: -M -p). Differential expression analysis was performed using DESeq2 v1.40.1. Pathway enrichment analysis was performed using the package pathfindR v2.2.0.

### Western blotting

1×10^6^ human peripheral blood mononuclear-derived macrophages per condition were treated with 1 μg/mL LPS, 20 ng/mL IFN-γ, 0.1 mM, 1 mM or 10 mM sodium butyrate, 10 μM Trichostatin A (TSA) (Sigma), 10 μM A485 (Selleck Chemicals), 10 μM RGFP966 (Sigma), and 10 μM TMP195 (Selleck Chemicals) for 24 hours. Then, cells were scraped and lysed using RIPA Lysis and Extraction Buffer (Thermo Scientific) with Protease inhibitor (Roche) and PhosSTOP (Roche). Lysates were quantified using Micro BCA Protein Assay Kit (Thermo Scientific). The same amount of lysate of each condition was loaded and run on 4-20% Mini-Protein TGX precast gels (Bio-Rad) at 100 V, and then blotted to PVDF membranes at 90 V, 60 minutes. Blots were blocked in 5% BSA (Fisher Scientific) at room temperature for 1 hour. After blocking, blots were washed three times in 0.01% TBST and then probed with polyclonal rabbit H3AcK27 antibody (Diagenode, C15410196) at 1:1000, polyclonal rabbit H3AcK4+9+14+18+23+27 (Ac-H3Kn) antibody at 1:1000 (Abcam, ab300641), monoclonal mouse Histone H3 antibody at 1:100 (Santa Cruz, sc-517576), polyclonal rabbit H4AcK5+8+12+16 (Ac-H4Kn) antibody at 1:1000 (Abcam, ab177790), anti-Acetyl-EP300-Lys1542 antibody at 1:500 (St ’ohn’s Laboratory, STJ97696), or monoclonal mouse GAPDH antibody at 1:1000 (Santa Cruz, sc-365062) at 4 °C overnight. The next day, blots were washed three times in 0.01% TBST and then labeled with Donkey anti Rabbit IgG IRDye680 (LI-COR, 926-32223) or Donkey anti Mouse IgG IRDye680 (LI-COR, 926-32222) both at 1:5000 at room temperature for 1 hour. After three times washing the blots were scanned using the Odyssey CLx Infrared Imaging System and analyzed using ImageStudio (LI-COR). Blots from the immunoprecipitation of EGFP-p300 were also probed with Ac-H3Kn antibody (Abcam, ab300641) and GAPDH antibody (Santa Cruz, sc-365062) both at 1:1000, polyclonal rabbit SP1 antibody (Proteintech, 21962-1-AP), and monoclonal mouse GFP (Roche, 11814460001) both at 1:500.

### Confocal microscopy

50,000 differentiated human macrophages per well were plated on 12-mm diameter glass coverslips and treated with 1 μg/mL LPS, 20 ng/mL IFN-γ, 0.1 mM, 1 mM or 10 mM sodium butyrate, 10 μM TSA, and 10 μM Thiomyristoyl for 24 hours. Cells were washed three times in PBS and fixed in pre-chilled methanol at –20°C for 10 minutes and blocked in PBS (Gibco) with 2% human serum (Sigma) at 4 °C for 1 hour followed by three times washing. Cells were incubated with primary antibody Ac-H3Kn (Abcam, ab300641) and Human Lamin A/C (BD, 612163) both at 1:500 in the blocking buffer at 4 °C overnight. The next day, cells were washed in PBS three times and coverslips were incubated for 1 hour at room temperature with the combination of the secondary antibodies Donkey anti-Rabbit IgG (H+L) Alexa Fluor 647 (Invitrogen, A31573) and Donkey anti-Mouse IgG (H+L) Alexa Fluor 488 (Invitrogen, A21202) both at 1:400. The coverslips were mounted in 70% glycerol with DAPI. Imaging was performed on a LSM800 Zeiss microscope with a 63× oil lens. Microscope images were analyzed using FIJI ImageJ. Pearson correlation coefficients (PCC) were calculated automatically using a macro in ImageJ (macro provided in Supplementary Information).

### Transfection and overexpression

1×10^7^ HeLa cells were plated in 10 mL DMEM medium in 75 cm^2^ cell culture flasks (Corning). eGFP-p300, eGFP-p300 (D1399Y), and eGFP-p300 (N1196D and R1234E) plasmids were transfected into HeLa cells using jetPEI DNA transfection reagent (Polyplus, 101000053). eGFP-p300 and eGFP-p300 (D1399Y) plasmids were gifts from Daniel Gerlich (Addgene plasmids #191764; #191763) (Schneider et al., 2022). eGFP-p300 (N1196D and R1234E) was generated from eGFP-p300 by point mutations. A mixture of 40 μg of plasmids and 160 μL jetPEI reagent was added to 1×107 HeLa cells and then incubated at 37 °C, 5% CO_2_ for 24 hours.

### Immunoprecipitation

HeLa cells were treated with 10 mM sodium butyrate, 10 μM A485 and 10 I-CBP112 (Sigma) for 24 hours, then cells were scraped and lysed using lysis buffer containing 25 mM Tris-HCl, 1% TritonX-100, 1 mM EDTA, 5% glycerol, and 150 mM NaCl, with Protease inhibitor and PhosSTOP. Lysates were quantified using the Micro BCA Protein Assay Kit (Thermo Scientific). 1 mg protein was combined with 2.4 μg polyclonal rabbit GFP antibody (Rockland, 600-401-215). The reaction volume was brought to 500 μL with the cell’s lysis buffer and incubated for 2 hours at room temperature. Subsequently, 60 μg washed Pierce Protein A/G Magnetic Beads (Thermo Scientific) were added per 2.4 μg GFP antibody. The mixture was incubated for 1 hour at room temperature. The beads were collected using a DynaMag magnets stand (Thermo Scientific) and washed in 0.05%TBST for three times. Finally, the beads were washed in UltraPure Distilled Water (Invitrogen), resuspended in 100 μL SDS-PAGE reducing sample buffer (Bio-Rad) and heated at 95°C for 10 minutes.

### MD simulations

The MD (Molecular Dynamics) simulations in this study involve four different protein systems related to the p300 acetyltransferase: Non-acetylated p300 based on structure PDB 6GYR (Ortega et al., 2018); Bromodomain, RING-loop and PHD-domains of non-acetylated p300 is based on structure PDB 6GYR, including residues 1045–1290; Bromodomain, RING-loop, and PHD-domain of p300 with three acetylated lysines based on structure PDB 6GYR with lysine residues 1180, 1203, 1228 converted to acetylated lysine; Bromodomain, RING-loop, and PHD-domain of p300 in contact with the histone protein of the nucleosome is based on structure PDB 8HAG (Kikuchi et al., 2023).

6GYR is a tetrameric structure; we have considered a single monomer. Additionally, the missing residues 1534–1565 from the HAT were reconstructed using MODELLER (Webb and Sali, 2016). To speed up the simulations, since we are interested in studying the interaction between the bromodomain and RING-loop, we built a smaller model comprising of the Bromodomain, RING-loop, and PHD-domains covering residues 1045–1290. Here, besides the non-acetylated form, we considered the triple acetylated form K1180Ac, K1203Ac, and K1128Ac where the three lysines in the RING-loop were converted to acetylated lysines. The parameters for lysine and the acetyl group are available in common force fields; however, a set of parameters (charges) specifically designed for the neutral group of acetylated lysine was missing. Therefore, we parameterized them to be compatible with the amber force field. We performed a QM calculation with the Gaussian16 program (Frisch et al., 2016) to generate the electrostatic potential, which was then used for deriving the RESP charges of the acetylated lysine (ACL) residues via Antechamber (Wang et al., 2006). After manually transformed the 1180, 1203 and 1230 residue from lysine to acetylated lysine, the structures were processed by the GROMACS protein builder gmx pdb2gmx using the all-atom AMBER model amber99sb-ildn (Lindorff-Larsen et al., 2010). A residue for the acetylated lysine was added to the library for the builder (abbreviated as ACL) with the new parameters. To study the interaction between the bromo-domain and the peptide chain of the histone protein, we constructed a reduced model based on the PDB 8HAG structure (Kikuchi et al., 2023). This structure contains the entire P300 acetyltransferase in contact with the nucleosome. Thus, as done previously, we kept only the bromodomain, RING-loop, and PHD-domain of the enzyme. Meanwhile, we kept only the first hundred residues of the final peptide chain (in contact with the bromodomain) of the histone protein of the nucleosome. This latter contains ACL residues, so here too we used the acetylated lysine parameters parameterized earlier. The protein systems were solvated TIP3P water molecules, adding Na^+^ and Cl^-^ ions to maintain neutrality and a salt concentration of 0.15 M. The amber99sb-ildn force field was used (Lindorff-Larsen et al., 2010). The velocity rescale thermostat (Bussi et al., 2007) was used to maintain the temperature at 300 K and an isotropic Parrinello–Rahman barostat (Parrinello and Rahman, 1981) with a reference pressure of 1 atm and isothermal compressibility of 4.5 × 10^-5^ bar^-1^ was used to maintain the pressure of the system. Long-range interactions were evaluated using particle-mesh Ewald summation (Darden et al., 1993). Following minimization and equilibration steps, the systems were simulated for 1 μs, using a time step of 2 fs. All the simulations were performed using the GROMACS-2021.5 package (Abraham et al., 2015).

### Enzyme-linked immunosorbent assay (ELISA)

PBMC-differentiated macrophages were seeded in the flat-bottom 96-well plate (Corning, 3596) at 100,000 cells/well, cells were treated with or without 1 μg/mL LPS, 20 ng/mL IFN-γ, sodium butyrate, and/or other reagents for 24 hours. TNF-α ELISA was performed using the ELISA kits according to the company protocols (Invitrogen, 88-7346).

## Data availability

ChIP-seq data have been deposited at the Gene Expression Omnibus (GEO) genomics data repository with accession number GSE255090. Plasmids that were generated for this study have been deposited at Addgene.

## Statistical analysis

Statistical analysis was performed on GraphPad software. Data were analyzed with paired 2 sides Student’s t-test for pairing of donors (mRNA, Western blot of human macrophages, confocal microscopy), or unpaired 2 sides Student’s t-test for immunoprecipitation of HeLa cells. The significance level was indicated by a P-value, as defined in the figure legends.

## Acknowledgements

We thank the Center for Information Technology of the University of Groningen for their support and for providing access to the Hábrók high performance computing cluster, and Prof. Dr. Daniel Gerlich from Vienna BioCenter of the Institute of Molecular Biotechnology of the Austrian Academy of Sciences for plasmids.

## Author contributions

Jiang, M. did the macrophage preparation for ChIP-seq, RT-qPCR, Western blot, and confocal microscopy; Chiariello, M. G. and Marrink, S.-J. performed the MD simulations; Modderman, R. and Jonkers, I. H. did the ChIP-seq; Incarnato, D. analysed the ChIP-seq data; Psoma, A. and Lyu, J. contributed to the mutagenesis, microscopy and ELISA experiments. Jiang, M. analysed data and wrote original draft with van den Bogaart, G.; Jiang, M., van den Bogaart, G., and Bianchi, F. conceptualized the study. All authors commented on the final draft.

