## Supplementary Information for "Butyrate blocks specific histone acetylation by preventing recruitment of p300 to acetylated histones"

**Supplementary Fig. 1**

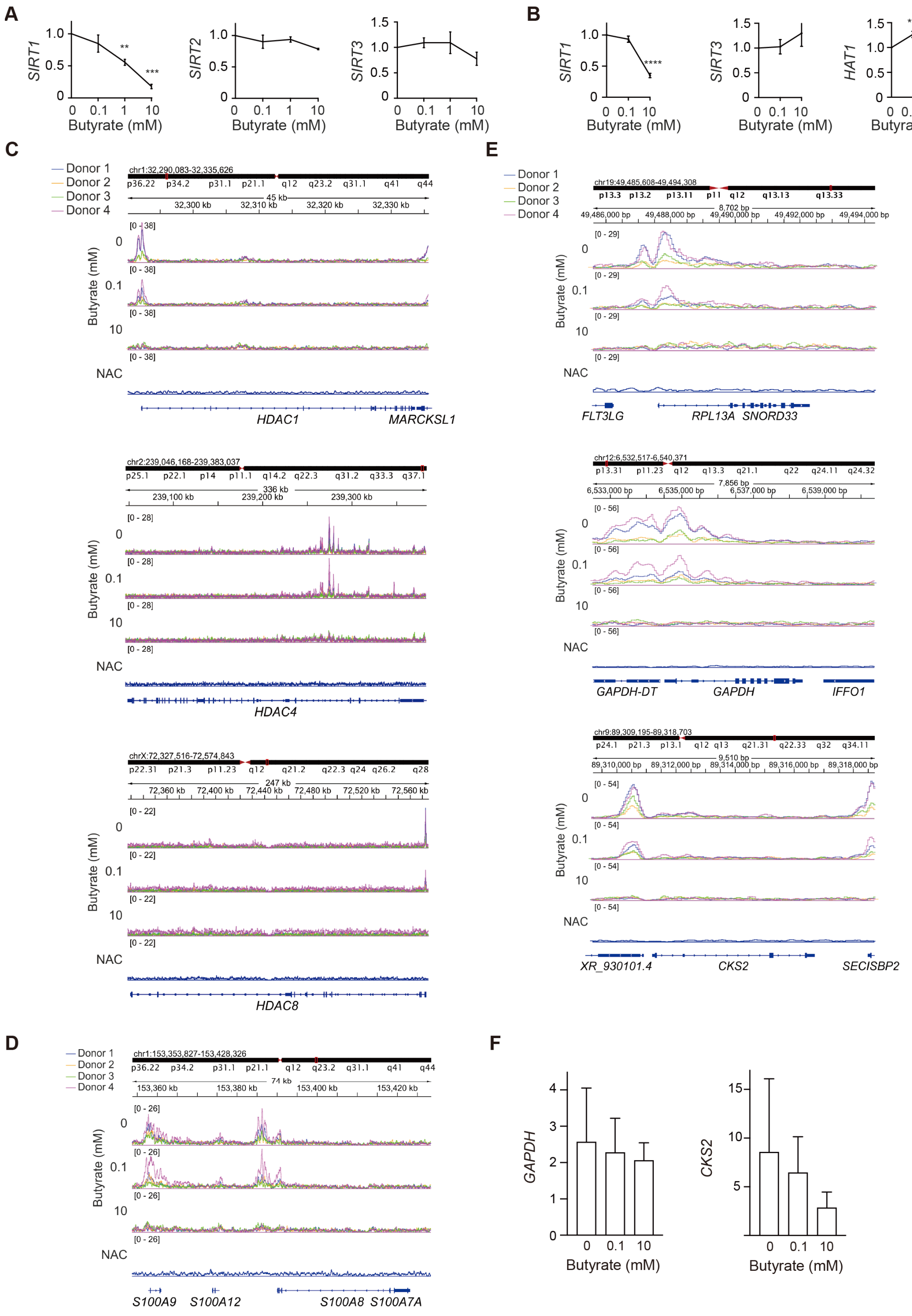


**Supplementary Figure 1.** RT-qPCR and ChIP-Seq of H3K27Ac. Human peripheral blood monocyte-derived macrophages were stimulated for 24 hr with LPS, IFN-γ and the indicated concentrations of Na-butyrate. (**A**) *SIRT1*, *SIRT2*, and *SIRT3* mRNA levels by RT-qPCR normalized to without butyrate. (One-way ANOVA with a Dunnett's multiple comparisons test, n=3 donors; error bars represent means ± SEM; **: P < 0.01; ***: P < 0.001; conditions were compared with the no butyrate condition) (**B**) *SIRT1*, *SIRT3*, and *HAT1* mRNA levels by RNA-seq normalized to human genome (One-way ANOVA with a Dunnett's multiple comparisons test; n=4 donors; error bars represent means ± SEM; ****: P < 0.0001). (**C-F**) Genome browser snapshots of published ChIP-seq data (Jiang et al., 2025) showing H3K27Ac enrichment in (**C**) *HDAC1, HDAC8, HDAC4,* (**D**) *S100A9, S100A8*, (**E**) *RPL13A, GAPDH,* and flanking genes. NAC: no antibody control. (**F**) *GAPDH* and *CKS2* mRNA levels by ChIP-qPCR normalized to genomic DNA (n = 4 donors; error bars represent means ± SEM)

**Supplementary Fig. 2**

**A**

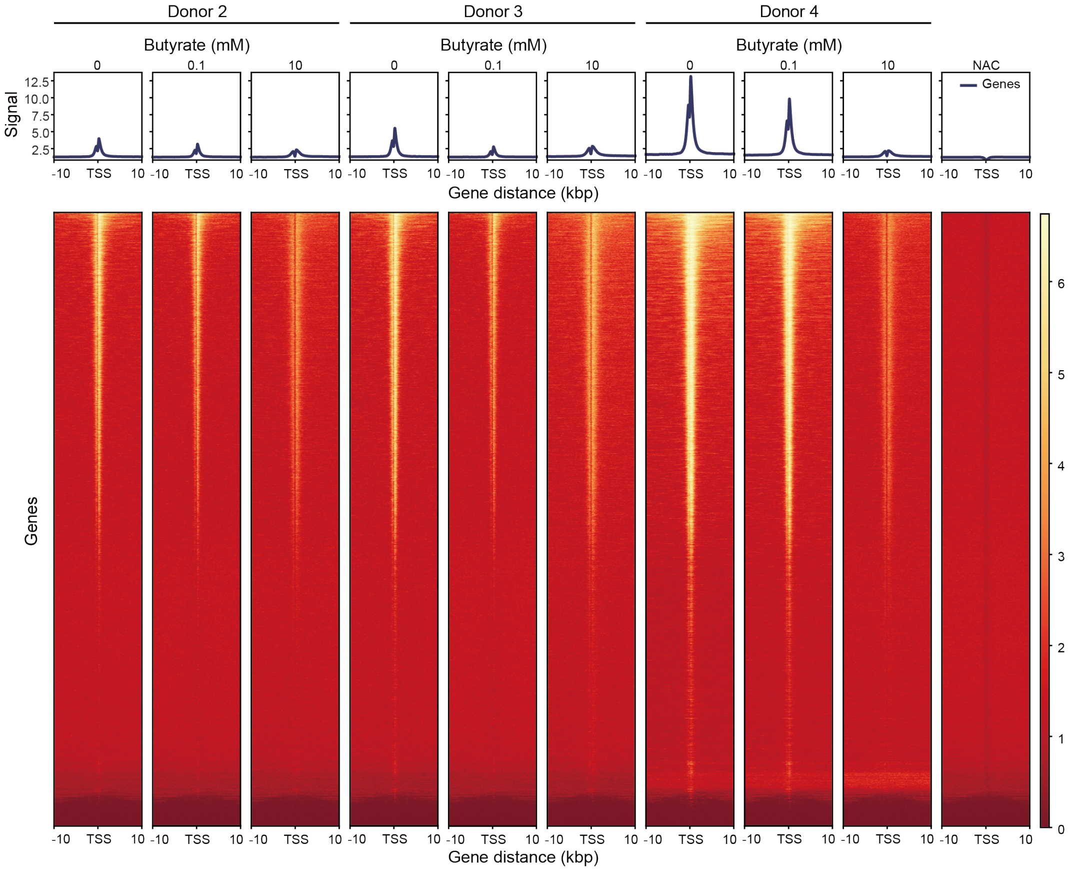

**B**

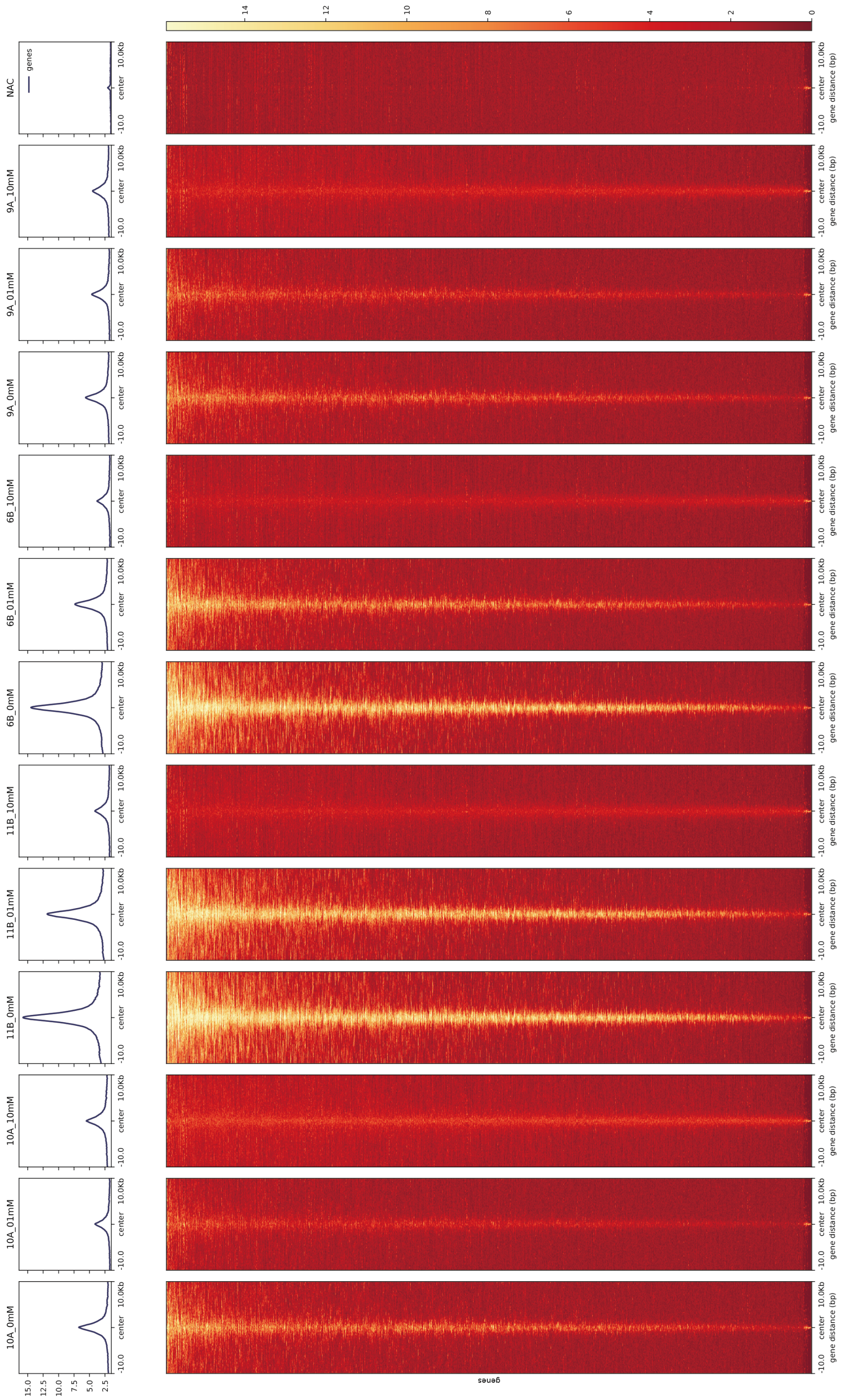

NAC
9A_10mM
9A_01mM
9A_0mM
6B_10mM
6B_01mM
6B_0mM
11B_10mM
11B_01mM
11B_0mM
10A_10mM
10A_01mM
10A_0mM
genes
-10.0
center
10.0Kb
gene distance (bp)
15.0
12.5
10.0
7.5
5.0
2.5
14
12
10
8
6
4
2
0


1

c

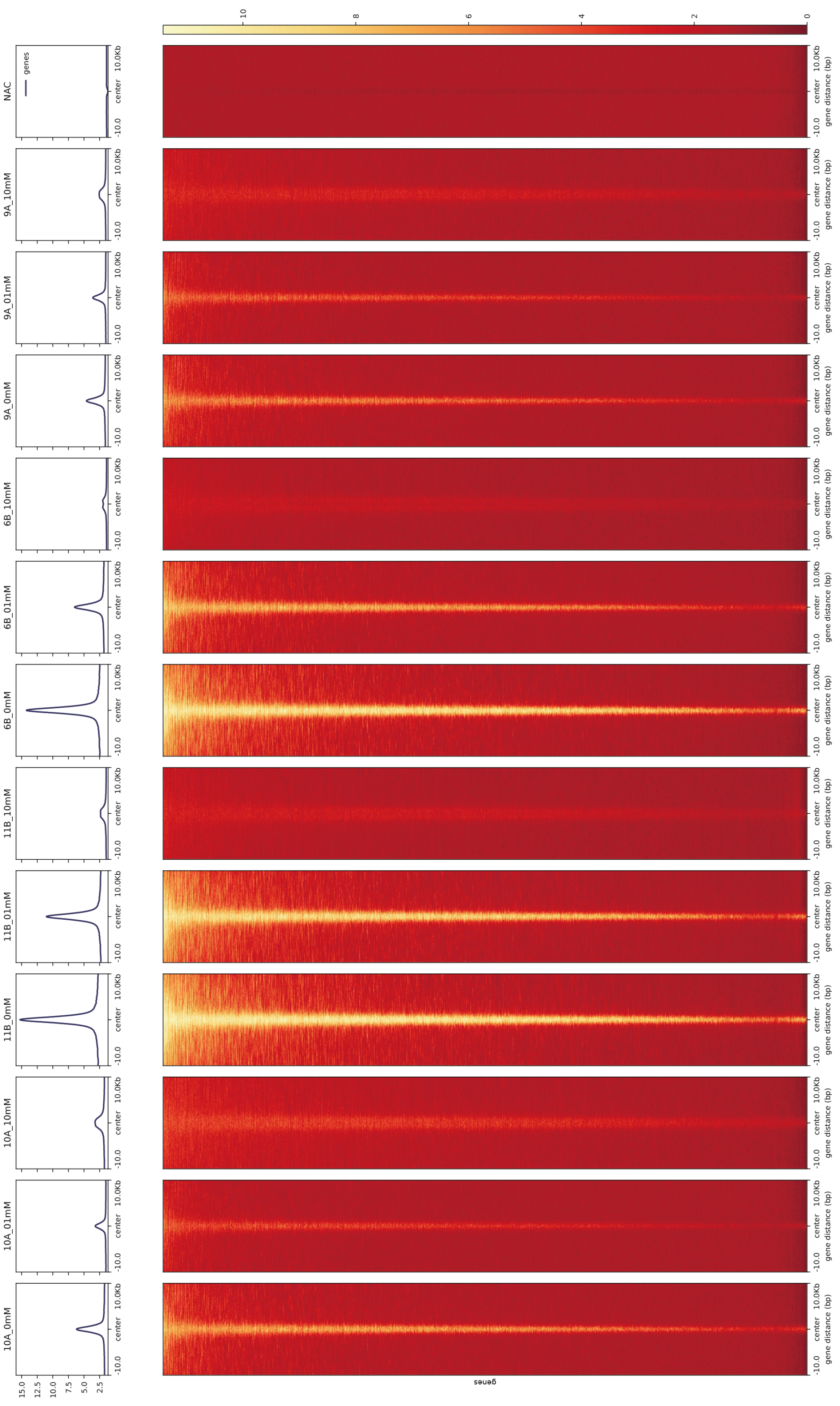

NAC
9A_10mM
9A_01mM
9A_0mM
6B_10mM
6B_01mM
6B_0mM
11B_10mM
11B_01mM
11B_0mM
10A_10mM
10A_01mM
10A_0mM
genes
-10.0
center
10.0Kb
gene distance (bp)
15.0
12.5
10.0
7.5
5.0
2.5
10
8
6
4
2
0


1

**D**

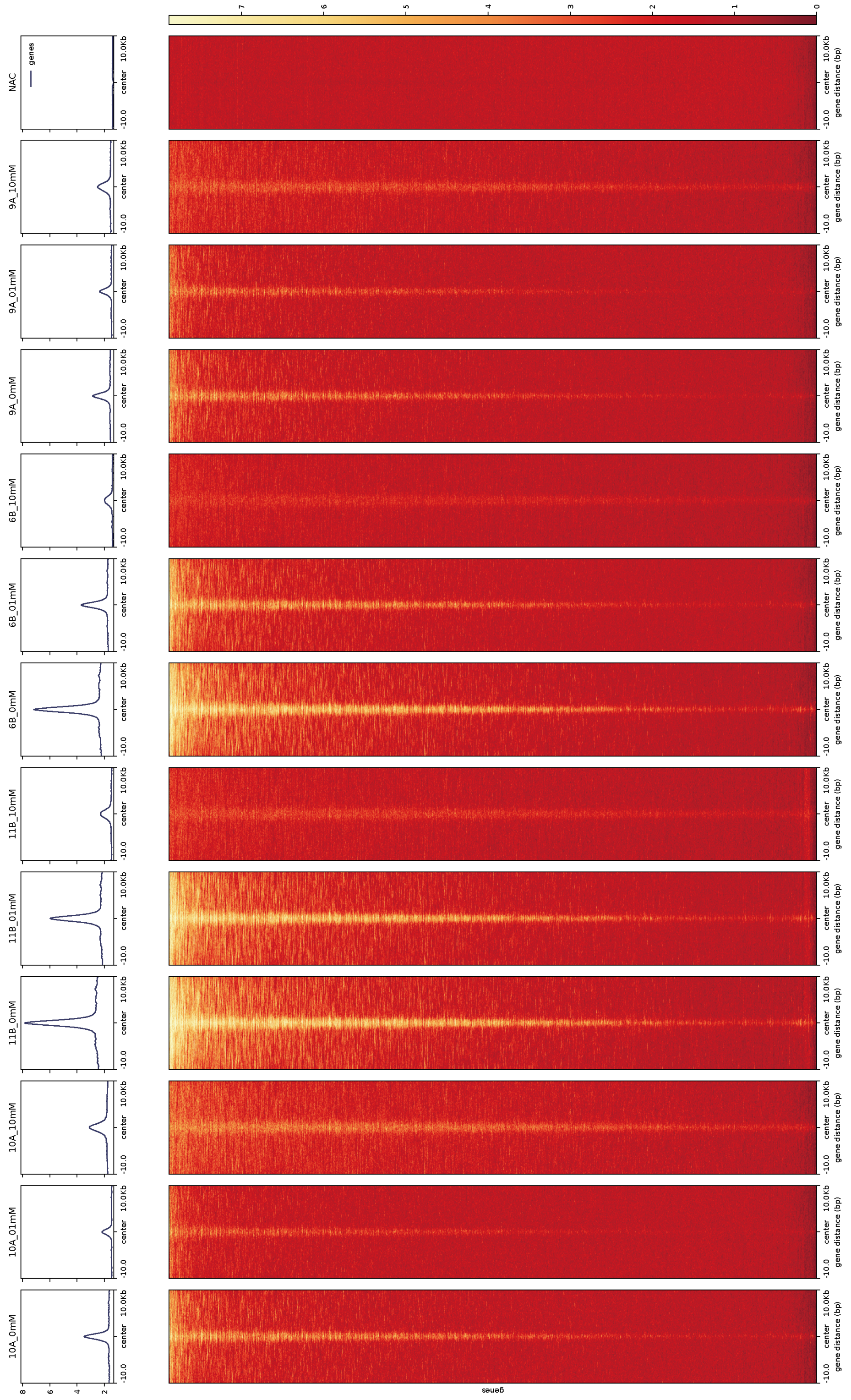


**Supplementary Figure 2.** Butyrate decreases histone acetylation at specific sites. (**A**) Intensity profile overlaps from published ChIP-seq data (Jiang et al., 2025) of all transcription start sites (TSS) for all 4 tested donors, showing a dose-dependent decrease in fold-enrichment of peaks relative to total reads. (**B**) Heatmap displays the subset of peaks, identified by peak calling of H3K27ac (using MACS 1.4.2), which are common to all 4 individuals at no butyrate condition. (**C**) Heatmap displays regions identified as significantly changing their acetylation level between 0 mM and 10 mM butyrate. These were identified by binning the genome using a sliding window of 1000 bp and by then using DeSeq2 (R) to identify regions of significant change in acetyl level (|log2(Fold-change)| >= 1, adjusted p-value < 0.05). (**D**) Heatmap displays regions corresponding to active enhancers in CD14-positive monocytes using ENCODE data. These were identified by selecting genomic regions that were H3K4me1 and H3K27ac positive, but H3K4me3 negative. The procedure was to first call peaks for H3K4me1, H3K4me3 and H3K27ac ChIP-seqs (using MACS 1.4.2), and then, using the intersect utility from BEDtools v2.31.0, overlapping H3K4me1 and H3K27ac peaks were selected, and any peak overlapping with a H3K4me3 peak was discarded.

**Supplementary Fig. 3**

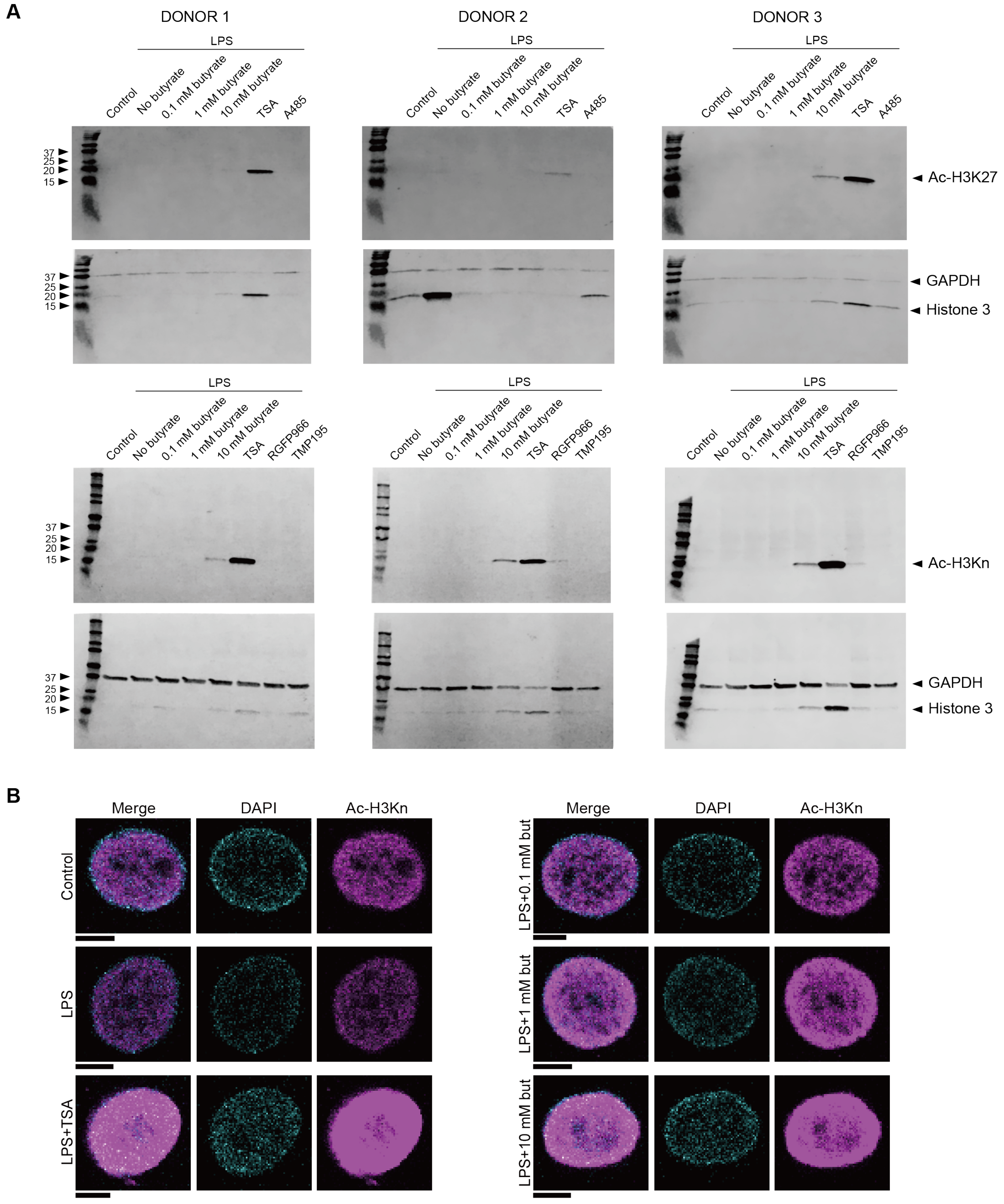


**Supplementary Figure 3.** Butyrate induces hyperacetylation of histones globally. Human peripheral blood monocyte-derived macrophages were stimulated for 24 hr with LPS, IFN-$\gamma$, the indicated concentrations of Na-butyrate, 10 $\mu$M HDAC inhibitor TSA, RGFP966, TMP195, and/or 10 $\mu$M HAT inhibitor A485. (**A**) Complete Western blots for 3 donors showing histone acetylation detected with two different antibodies recognizing acetylated (Ac-)H3K27 and H3K4+9+14+18+23+27 (Ac-H3Kn). Blots were stripped and reprobed for GAPDH (loading control) and total H3, however, the total H3 showed poor staining possibly due to incomplete stripping, epitope availability and/or antibody quality. (**B**) Representative confocal images from 3 donors for histone acetylation (Ac-H3Kn; magenta) and DAPI (cyan) (at least 5 cells per donor). Some pictures of the Ac-H3Kn staining are saturated to highlight the increase in staining intensity. Scale bars: 5 µm.

**Supplementary Fig. 4**

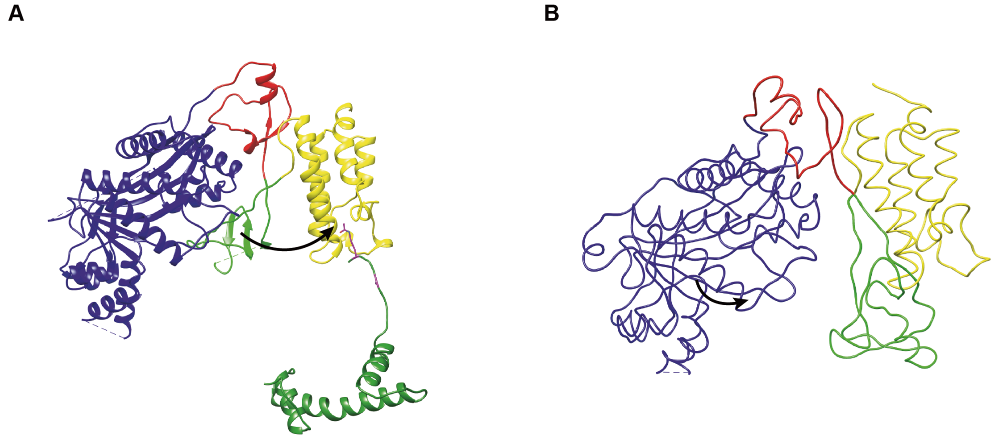


**Supplementary Figure 4.** Comparison of cryo-EM structures suggests that the RING loop moves between the HAT and bromodomain. Different protein sub-units are color-coded: bromodomain (yellow), RING-loop (green), PHD-domain (red), HAT-domain (blue), and histone H4 (dark green). (**A**) Cryo-EM model of p300, with its Bd bound to the acetylated histone 4 (H4) of the H4K12acK16ac nucleosome (pdb code 8HAG) (Kikuchi et al., 2023). For clarity, only p300 and acetylated H4 is shown. (**B**) Low resolution cryo-EM structure of p300 without substrate (pdb code: 6K4N) (Ghosh et al., 2019). Note that the RING loop is tilted towards the binding pocked of the Bd (arrows).

**Supplementary Fig. 5**

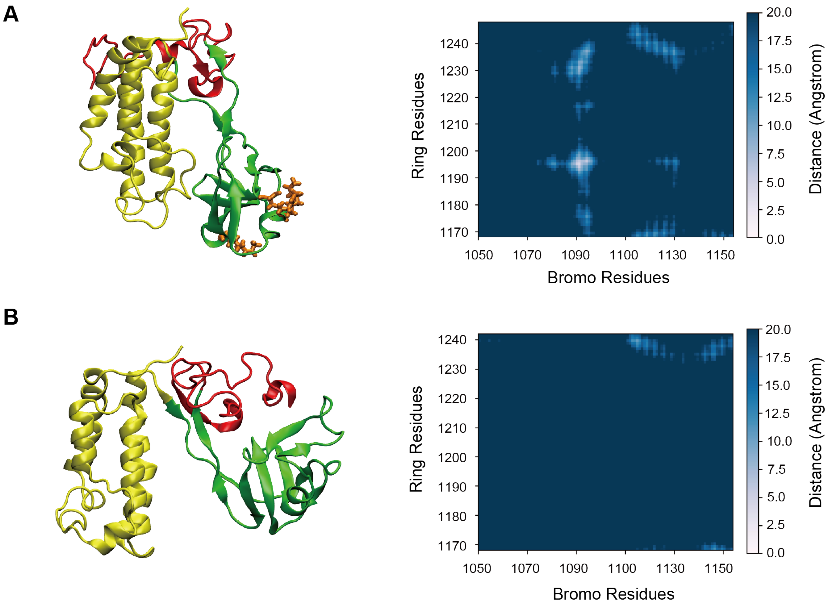


**Supplementary Figure 5.** MD simulations of the p300 acetyltransferase system. The reference structure is PDB 6GYR. Different protein sub-units are color-coded: bromodomain (yellow), RING-loop (green), and PHD-domain (red). Structure extracted from the simulation of the triple acetylated form K1180Ac, K1203Ac, and K1228Ac, with acetylated lysines highlighted in orange (**A**) and the non-acetylated form (**B**). The contact maps between the bromodomain and RING-loop residues are shown in the right panel for both the systems.

**Supplementary Fig. 6**

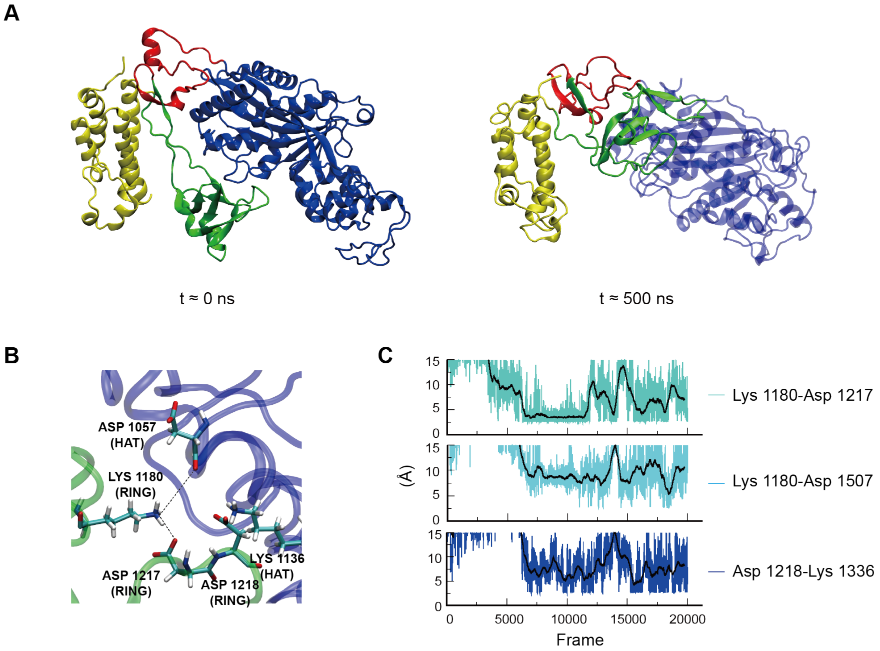


**Supplementary Figure 6.** MD simulations of the full non-acetylated p300 acetyltransferase system. Different protein sub-units are color-coded: bromodomain (yellow), RING-loop (green), PHD-domain (red) and HAT (blue). (**A**) Representative structures at time 0 and 500 ns simulated time. (**B**) The RING-loop is attached to the HAT through a network of interactions involving K1180. (**C**) H-bond interactions between the RING-loop and HAT over the simulation time.

**Supplementary Fig. 7**

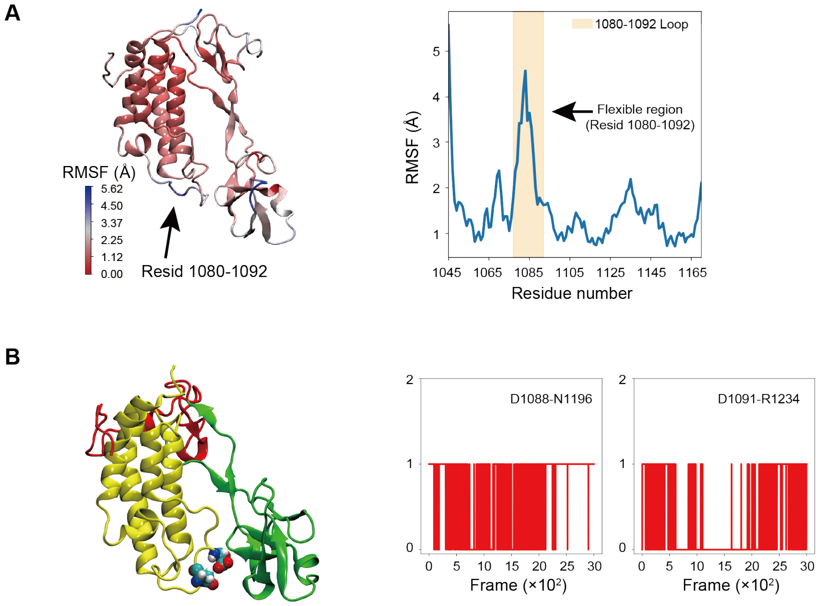

**Supplementary Figure 7.** MD simulations of the interactions between the bromodomain and RING-loop. (**A**) Root mean square fluctuations (RMSF) of the bromodomain (residue number 1045–1168). The structure colored according to the RMSF value is also shown. (**B**) Number of contacts between D1088 and D1091 (bromodomain) and N1196 and R1234 (RING-loop) over the simulation time. Representative structure with the D1088 and D1091 in ball-and-stick representation located on the flexible loop of the bromodomain.

**Supplementary Fig. 8**

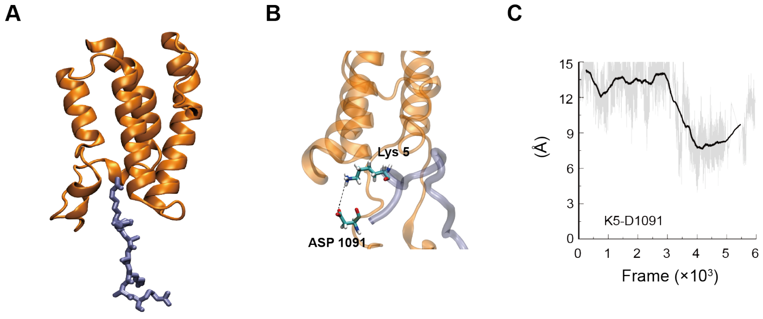


**Supplementary Figure 8.** MD simulations of the p300 acetyltransferase system bound to acetylated histone H4. The reference structure is PDB 8HAG. Different protein sub-units are color-coded: bromodomain (orange), histone H4 N-terminal peptide chain (ice blue). (**A**) Histone chain buried into the bromo domain from the cryo EM structure 8HAG. Here the residues 1–11 are missing. (**B**) Snapshot from MD simulation, where the missing residues 1–11 were reconstructed. The interaction between K5 and D1091 is highlighted. (**C**) The distance between K5 and D1091 over the simulation frames is shown.

**Supplementary Fig. 9**

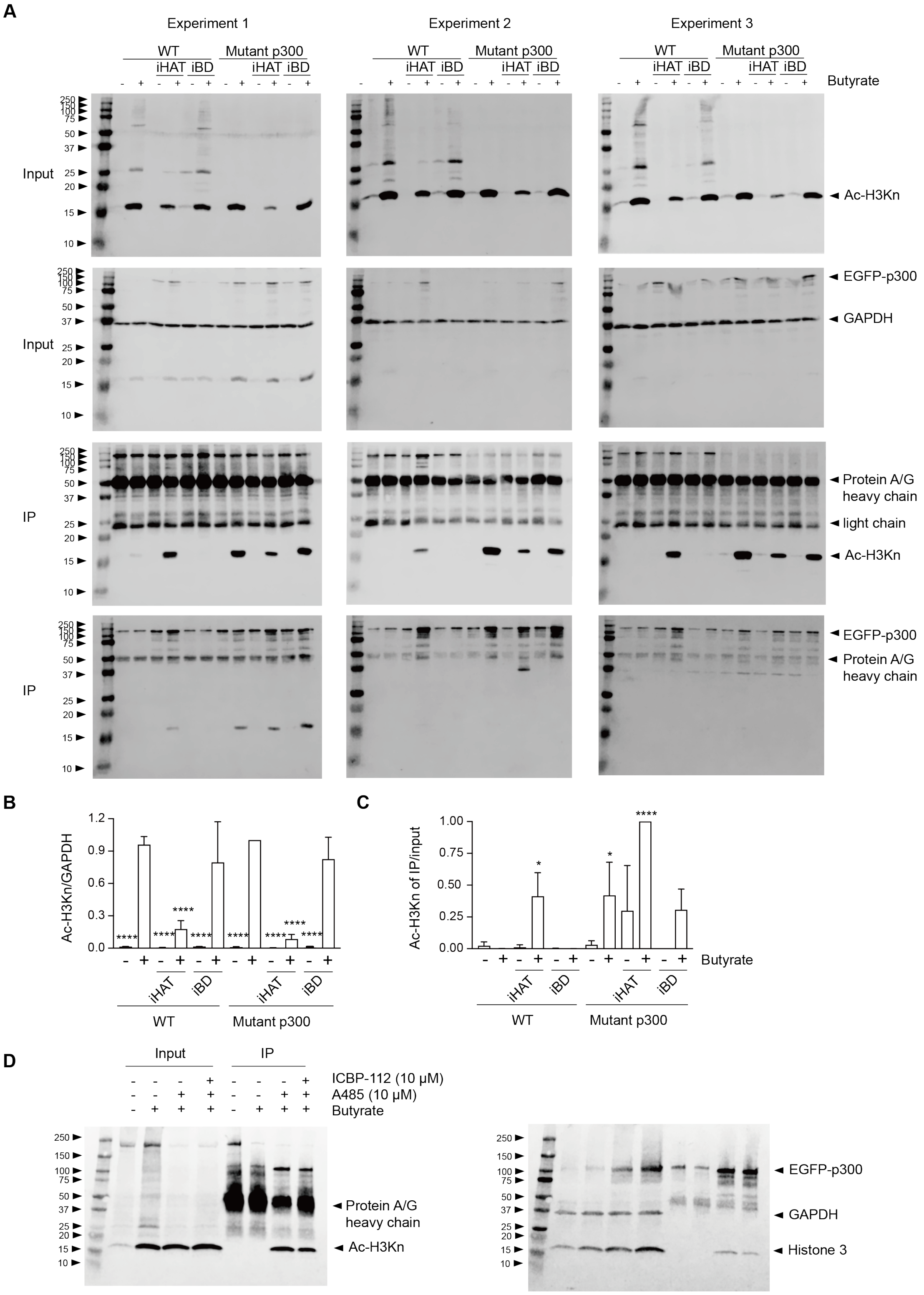

**E**

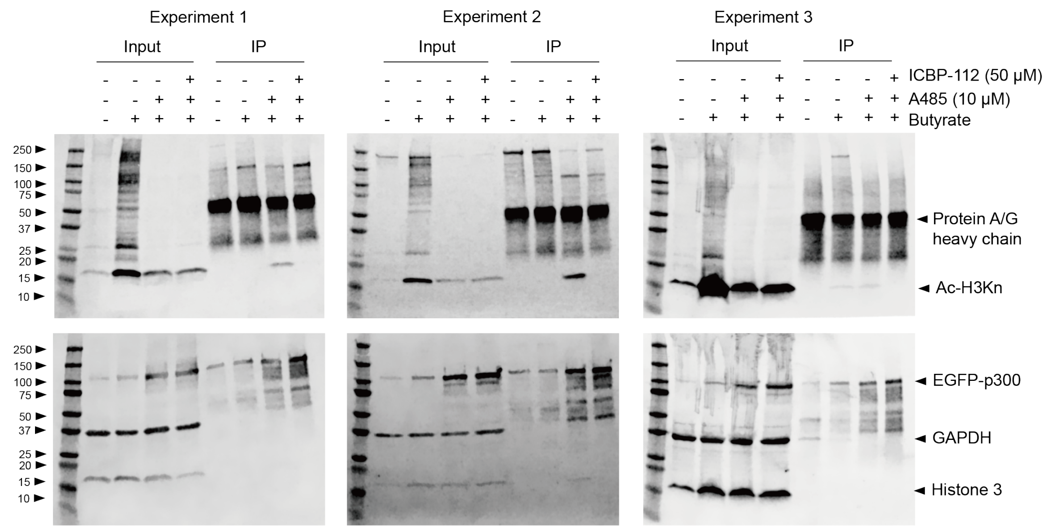
Experiment 1
Experiment 2
Experiment 3
Input
IP
ICBP-112 (50 μM)
A485 (10 μM)
Butyrate
250
150
100
75
50
37
25
20
15
10
Protein A/G
heavy chain
Ac-H3Kn
EGFP-p300
GAPDH
Histone 3

**F**

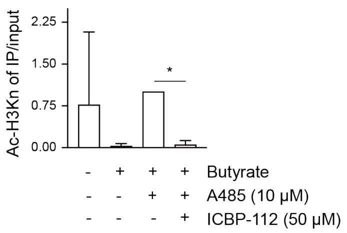
Ac-H3Kn of IP/input
2.25
1.50
0.75
0.00
*
Butyrate
A485 (10 μM)
ICBP-112 (50 μM)

**G**

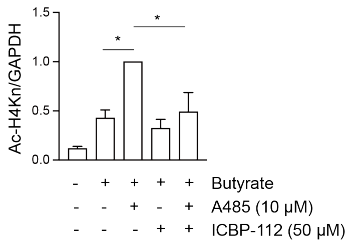
Ac-H4Kn/GAPDH
1.5
1.0
0.5
0.0
*
*
Butyrate
A485 (10 μM)
ICBP-112 (50 μM)

**H**

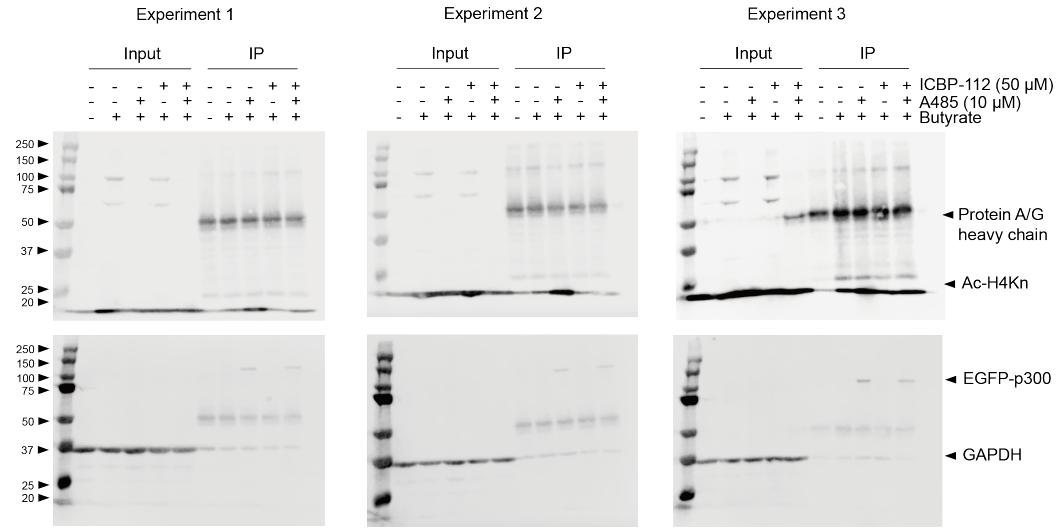
Experiment 1
Experiment 2
Experiment 3
Input
IP
ICBP-112 (50 μM)
A485 (10 μM)
Butyrate
250
150
100
75
50
37
25
20
Protein A/G
heavy chain
Ac-H4Kn
EGFP-p300
GAPDH

**I**

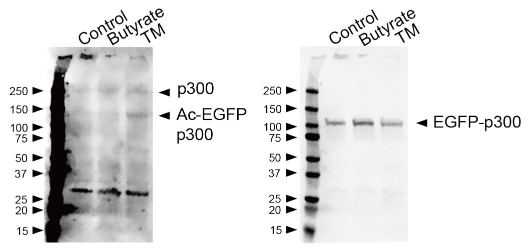
Control
Butyrate
TM
250
150
100
75
50
37
25
20
15
p300
Ac-EGFP
p300
EGFP-p300

1

**J**

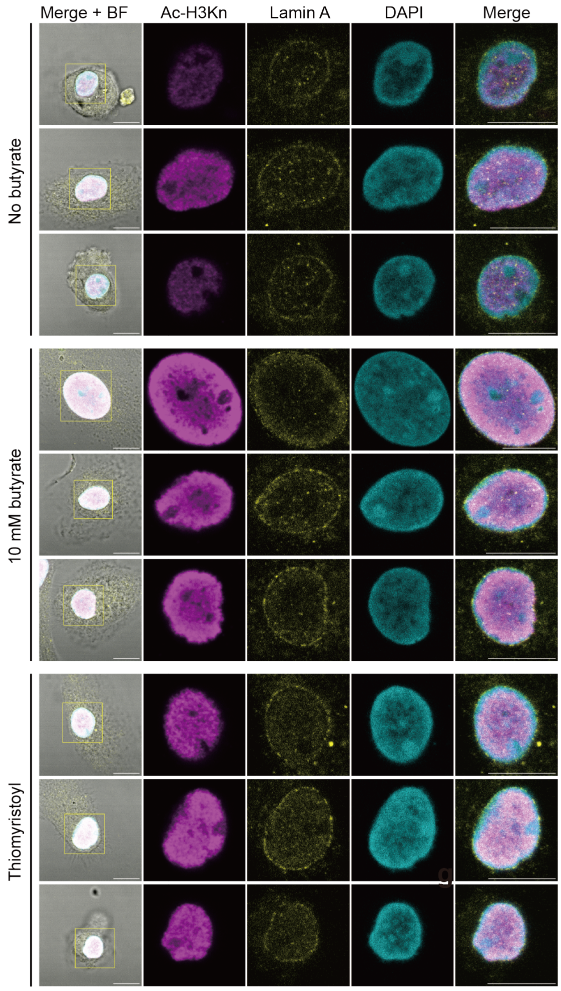


**Supplementary Figure 9.** Butyrylated p300 does not bind to acetylated H3. (**A**) HeLa cells were transfected with EGFP-tagged catalytic core of p300 wildtype or catalytically inactive D1399Y mutant and stimulated for 24 hr with 10 mM Na-butyrate, 10 μM HAT inhibitor, and/or 10 μM bromodomain inhibitor prior to IP for EGFP. Complete Western blots for 3 independent experiments, showing Ac-H3K4+9+14+18+23+27 (Ac-H3Kn), EGFP-p300, and GAPDH detected in lysates and IP samples. (**B**) Quantification of Ac-H3Kn levels normalized to GAPDH in lysates. (**C**) Quantification of Ac-H3Kn levels in IP samples normalized to lysates (One-way ANOVA with a Dunnett's multiple comparisons test; n=3 independent experiments; error bars represent means ± SEM; *: $P < 0.05$; ****: $P < 0.0001$; each condition was compared to the butyrate-treated WT condition). (**D** and **E)** Transfected Hela cells were stimulated for 24 hr with 10 mM Na-butyrate, 10 μM A485, and 10 μM (**D**) or 50 μM (**E**) ICBP-112. (**F**) Quantification of

Ac-H3Kn levels in IP samples normalized to lysates (One-way ANOVA with a Tukey's multiple comparisons test; n = 3 independent experiments, error bars represent means ± SEM. *: $P < 0.05$). (**G**) Quantification of Ac-H4Kn levels in IP samples normalized to GAPDH in lysates (One-way ANOVA with a Tukey's multiple comparisons test; n = 3 independent experiments, error bars represent means ± SEM. *: $P < 0.05$). (**H**) Transfected Hela cells were stimulated for 24 hr with 10 mM Na-butyrate, 10 μM A485, and 50 μM ICBP-112. Western blot showing Ac-H4K5+8+12+16 (Ac-H4Kn) in lysates and IP samples. (**I**) Transfected Hela cells were stimulated for 24 hr with 10 mM Na-butyrate, 10 μM of SIRT2 inhibitor Thiomyristoyl (TM), or 10 μM A485 followed by Western blot with an antibody recognizing an acetylated lysine located in the AIL of p300 (K1542). The experiment shows that acetylated p300 was increased by TM treatment. (**J**) Human peripheral blood monocyte-derived macrophages were stimulated for 24 hr with LPS, IFN-γ, and the indicated concentrations of Na-butyrate or 10 μM TM. Representative confocal images from 3 donors for Ac-H3Kn (magenta) (at least 5 cells per donor), Lamin A (yellow), and DAPI (cyan). Scale bars: 5 μm.

**Supplementary Fig. 10**

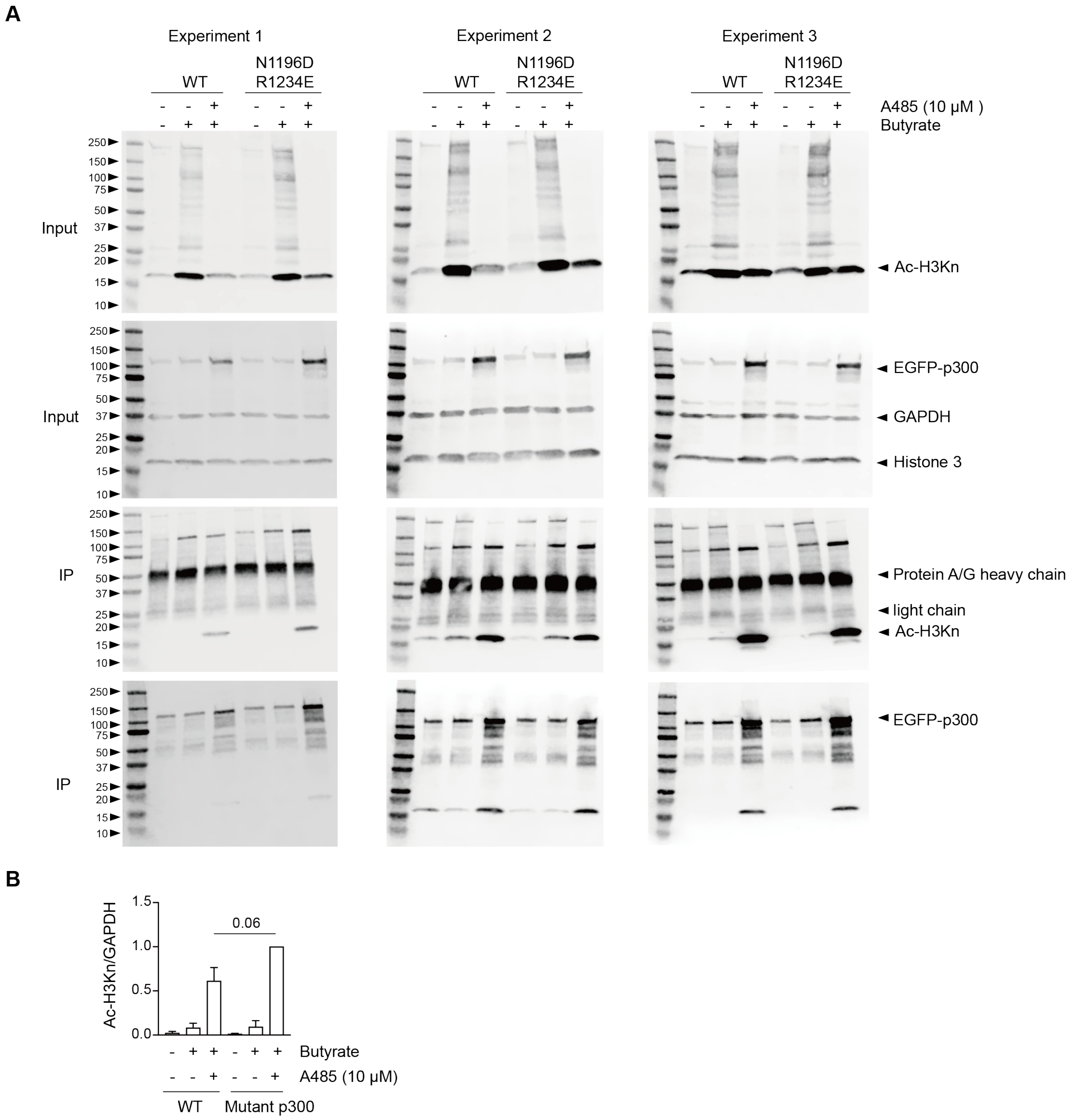


**Supplementary Figure 10.** N1196 and R1234 of RING loop suppress p300 bromodomain binding to acetylated histone. (**A**) Hela cells were transfected with the EGFP tagged catalytic core of p300 wildtype or N1196D and R1234E mutant, and stimulated for 24 hr with 10 mM Na-butyrate, 10 μM HAT inhibitor A485 to immunoprecipitation (IP) for EGFP. Western blot showing Ac-H3K4+9+14+18+23+27 (Ac-H3Kn), EGFP-p300 and GAPDH detected in lysates and IP samples. (**B**) Quantification of Ac-H3Kn levels normalized to GAPDH in lysates (One-way ANOVA with a Tukey's multiple comparisons test; n = 3 independent experiments, error bars represent means ± SEM. *: $P < 0.05$)

**Supplementary Fig. 11**

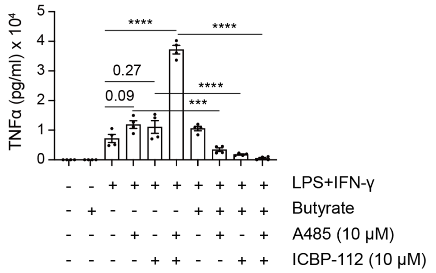


**Supplementary Figure 11.** Opposite effects of p300 inhibition on TNF-α production by macrophages in the presence and absence of butyrate. Human PBMC-derived macrophages were stimulated for 24 hr with LPS, IFN-γ, 10 mM Na-butyrate, 10 μM HAT inhibitor A485, and/or 10 μM bromodomain inhibitor ICBP-112. TNF-α production was determined by ELISA (One-way ANOVA with a Tukey's multiple comparisons test; n=4 donors; error bars represent means ± SEM; ***: $P < 0.001$; ****: $P < 0.0001$).

**Supplementary Fig. 12**

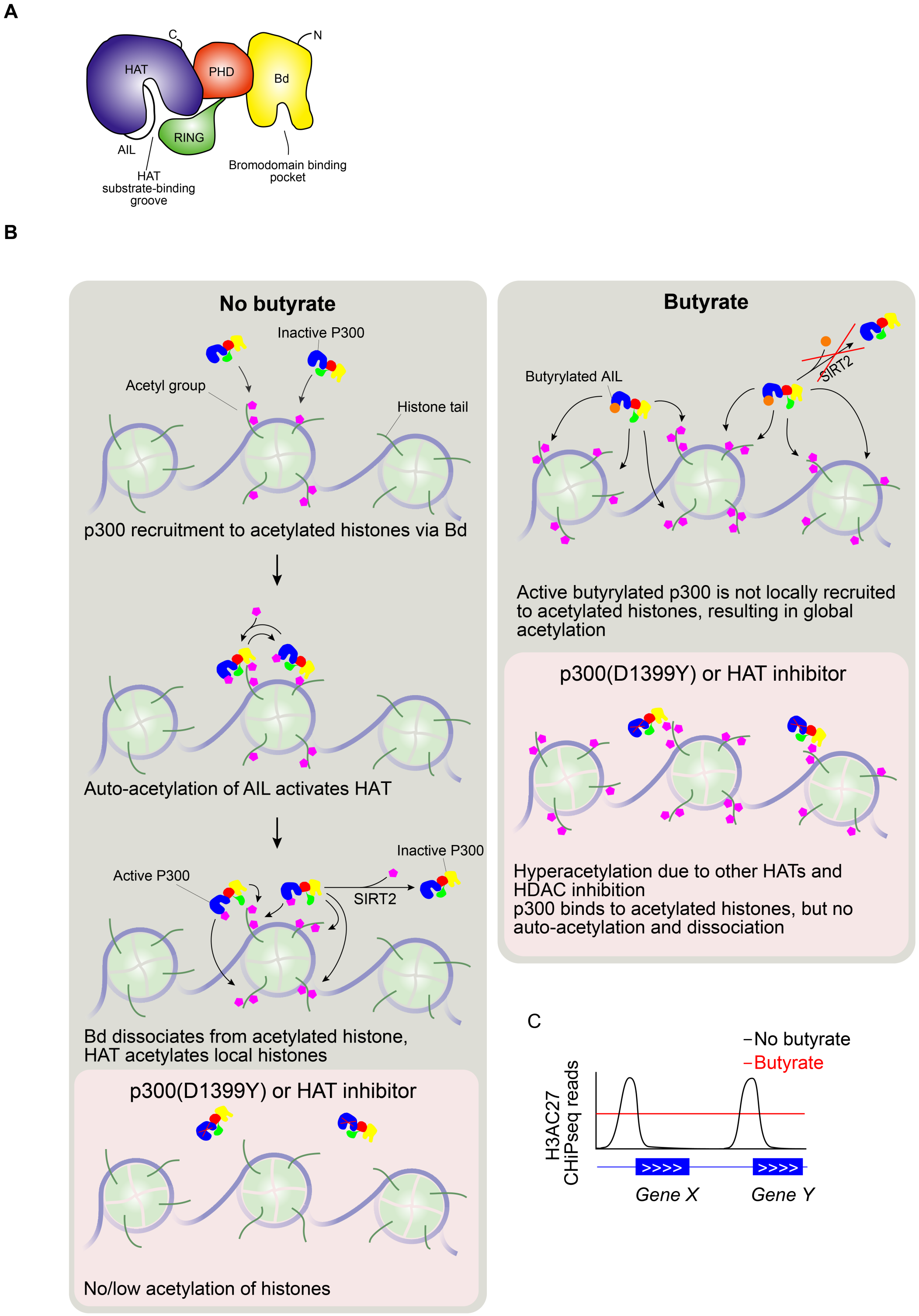


**Supplementary Figure 12.** Model for loss of specific histone acetylation by butyrate. (**A**) Domain topology of the catalytic core of p300. Yellow: bromodomain (Bd); green: RING-loop: red, PHD domain: blue, HAT domain: AIL: autoinhibitory loop. Adapted from (Ortega et al., 2018). (**B**) Proposed model for histone acetylation by p300 in the absence and presence of butyrate. In the absence of butyrate, active p300 molecules are recruited to acetylated histones by binding to its Bd. This promotes in-*trans* auto-acetylation of the AIL, triggering its activation. Lysines located in the RING-loop also become acetylated, weakening the interaction with the HAT. The RING-loop can now interact with the bromodomain and thereby prevent binding to acetylated histones. Activated p300 dissociates from acetylated histones and acetylates other histones nearby. p300 is inactivated by SIRT2, which deacetylates the AIL. Histone acetylation is reduced with the catalytically inactive p300 mutant or with a HAT inhibitor. Butyrate induces butyrylation of the AIL of p300, resulting in its hyperactivation. However, the Bd of butyrylated p300 is not recruited to acetylated histones. In addition, SIRT2 is less effective in removing p300 butyrylation compared to the acetylation. These effects result in non-specific hyperacetylation of histones. Butyrylation of p300 is blocked with a catalytically inactive mutant or with a p300 inhibitor, and the Bd of p300 remains associated to acetylated histones. In these conditions, butyrate still increases histone acetylation due to its inhibition of HDACs and the activities of other HATs. (**C**) Butyrate results in global increase of histone acetylation, and loss of specific histone acetylation.

**Supplementary Table 1. Primer sequences for reverse transcription-quantitative PCR.**

| Gene | Primer sequence (5'-3') |
|---|---|
| ***HDAC1*** | F: GGTCCAAATGCAGGCGATTCCT<br>R: TCGGAGAACTCTTCCTCACAGG |
| ***HDAC3*** | F: GAGTTCTGCTCGCGTTACACAG<br>R: CGTTGACATAGCAGAAGCCAGAG |
| ***HDAC4*** | F: AGGTGAAGCAGGAGCCCATTGA<br>R: GGTAGTTCCTCAGCTGGTGGAT |
| ***HDAC7*** | F: TCCTGGCACAGCGGATGTTTGT<br>R: TGAAGGCGAGGTCAGTGACACT |
| ***HDAC8*** | F: TGTGCTGGAAATCACGCCAAGC<br>R: ACCACTCCTCAGCTCTGGAAAC |
| ***SIRT1*** | F: TAGACACGCTGGAACAGGTTGC<br>R: CTCCTCGTACAGCTTCACAGTC |
| ***SIRT2*** | F: CTGCGGAACTTATTCTCCCAGAC<br>R: CCACCAAACAGATGACTCTGCG |
| ***SIRT3*** | F: CCCTGGAAACTACAAGCCCAAC<br>R: GCAGAGGCAAAGGTTCCATGAG |
| ***EP300*** | F: GATGACCCTTCCCAGCCTCAAA<br>R: GCCAGATGATCTCATGGTGAAGG |
| ***SNRPD3*** | F: GGAAGCTCATTGAAGCAGAGGAC<br>R: CAGAAAGCGGATTTTGCTGCCAC |

HDAC, histone deacetylase; SIRT, Sirtuin; SNRPD3, Small Nuclear Ribonucleoprotein D3 Polypeptide.

**Supplementary Data 1. FIJI macro for automated quantification of the mean**

**intensity of histone acetylation normalized to DAPI.**

```
var bittype = 16;
var directory = File.directory;
var id0 = getImageID();
var name0 = getTitle();
run("Duplicate...", "duplicate channels=1");
if (bittype == 8)
{setMinAndMax(1, 1); //8bit}
else {setMinAndMax(100, 100); //16-bit}
run("Apply LUT");
run("8-bit");
rename("ch1");
var id1 = getImageID();
var name1 = getTitle();
selectImage(id0);
if (bittype == 8)
{run("Duplicate...", "duplicate channels=3");//8-bitsetMinAndMax(1, 1); //8bit}
else {run("Duplicate...", "duplicate channels=4");//16-bitsetMinAndMax(100, 100); //16-bit}
```

```
run("Apply LUT");
run("8-bit");
rename("ch4");
var id2 = getImageID();
var name2 = getTitle();
imageCalculator("Add create", "ch1","ch4");
selectWindow("Result of ch1");
run("32-bit");
var id3 = getImageID();
setAutoThreshold("Default dark no-reset");
//run("Threshold...");
run("NaN Background");
setMinAndMax(0, 1);
selectImage(id1);
close();
selectImage(id2);
close();
selectImage(id0);
run("Duplicate...", "duplicate channels=1");
```

```
rename("ch1");
var id2 = getImageID();
var name2 = getTitle();
imageCalculator("Multiply create 32-bit", "Result of ch1","ch1");
selectWindow("Result of Result of ch1");
var id5 = getImageID();
run("Divide...", "value=255");
if (bittype == 16)
{run("Divide...", "value=256"); //16-bit}
run("Measure");
selectImage(id3);
close();
selectImage(id2);
close();
selectImage(id0);
close();
selectImage(id5);
close();
```

**Supplementary Data 2. FIJI macro for calculation of the Pearson correlation**

**coefficient (PCC) between the Lamin A and H3Ac staining.**

```
var ID1 = getImageID();
selectImage(ID1);
run("Duplicate...", "duplicate channels=1");
var ID1_ch1 = getImageID();
var Name1_ch1 = getTitle();
selectImage(ID1);
run("Duplicate...", "duplicate channels=2");
var ID1_ch2 = getImageID();
var Name1_ch2 = getTitle();
run("JACoP ", "imga=[" + Name1_ch1 + "] imgb=[" + Name1_ch2 + "] pearson");
selectImage(ID1_ch1);
close();
selectImage(ID1_ch2);
close();
selectImage(ID1);
close();
```
